# Structural basis for helicase-polymerase coupling in the SARS-CoV-2 replication-transcription complex

**DOI:** 10.1101/2020.07.08.194084

**Authors:** James Chen, Brandon Malone, Eliza Llewellyn, Michael Grasso, Patrick M. M. Shelton, Paul Dominic B. Olinares, Kashyap Maruthi, Ed Eng, Hasan Vatandaslar, Brian T. Chait, Tarun Kapoor, Seth A. Darst, Elizabeth A. Campbell

**Author notes:** These authors contributed equally: James Chen, Brandon Malone.

## Abstract

SARS-CoV-2 is the causative agent of the 2019-2020 pandemic. The SARS-CoV-2 genome is replicated-transcribed by the RNA-dependent RNA polymerase holoenzyme (subunits nsp7/nsp8_2_/nsp12) along with a cast of accessory factors. One of these factors is the nsp13 helicase. Both the holo-RdRp and nsp13 are essential for viral replication and are targets for treating the disease COVID-19. Here we present cryo-electron microscopic structures of the SARS-CoV-2 holo-RdRp with an RNA template-product in complex with two molecules of the nsp13 helicase. The Nidovirus-order-specific N-terminal domains of each nsp13 interact with the N-terminal extension of each copy of nsp8. One nsp13 also contacts the nsp12-thumb. The structure places the nucleic acid-binding ATPase domains of the helicase directly in front of the replicating-transcribing holo-RdRp, constraining models for nsp13 function. We also observe ADP-Mg^2+^ bound in the nsp12 N-terminal nidovirus RdRp-associated nucleotidyltransferase domain, detailing a new pocket for anti-viral therapeutic development.

## INTRODUCTION

Coronaviruses (CoVs) are positive-strand RNA (+RNA) viruses belonging to order *Nidovirales* (de Groot et al., 2012). These viruses are responsible for several zoonotic infections (Snijder et al., 2016). Deadly incidences include the 2003 Severe Acute Respiratory Syndrome (SARS) pandemic caused by SARS-CoV-1 and the 2012 and 2015 Middle Eastern Respiratory Syndrome (MERS) outbreak and pademic caused by MERS-CoV (Hilgenfeld and Peiris, 2013). SARS-CoV-2, a β-CoV, has been identified as the cause of the current globally devastating CoV Disease 2019 (COVID-19) pandemic (Wu et al., 2020; Zhou et al., 2020).

The CoV family of viruses contains the largest known RNA genomes (Nga et al., 2011). An RNA-dependent RNA polymerase (RdRp, encoded by non-structural protein, or nsp12) functions in a holo-RdRp (comprising nsp7/nsp8_2_/nsp12) to synthesize all viral RNA molecules (Kirchdoerfer and Ward, 2019; Subissi et al., 2014). The RdRp is the target for antivirals such as remdesivir (Agostini et al., 2018; Yin et al., 2020). In addition to its central role in replication and transcription of the viral genome, the RdRp contains an N-terminal nidovirus RdRp-associated nucleotidyltransferase (NiRAN) domain with unknown function but with enzymatic activity that is also essential for viral propagation (Lehmann et al., 2015a).

The holo-RdRp is thought to coordinate with a number of accessory factors during the viral life cycle (Snijder et al., 2016; Sola et al., 2015). One of these accessory factors is nsp13, a superfamily 1B (SF1B) helicase that can unwind DNA or RNA in an NTP-dependent manner with a 5’ -> 3’ polarity (Ivanov and Ziebuhr, 2004; Lee et al., 2010; Seybert et al., 2000; SEYBERT et al., 2000; Tanner et al., 2003). The helicase is essential for replication in the nidovirus equine arteritis virus (EAV) (Dinten et al., 2000; Seybert et al., 2000, 2005), and the β-CoV murine hepatitis virus (Zhang et al., 2015), and is presumed to be essential in all nidoviruses (Lehmann et al., 2015b). The helicase is thought to play crucial roles in many aspects of the viral life cycle, some of which can be uncoupled by point mutations (Dinten et al., 1996; Marle et al., 1999). Besides helicase activities, nsp13 harbors RNA 5’-triphosphatase activity that may play a role in mRNA capping (Ivanov and Ziebuhr, 2004; Ivanov et al., 2004).

In addition to the two canonical RecA ATPase domains of SF1 helicases (Saikrishnan et al., 2009; Singleton et al., 2007), nsp13 contains three domains unique to nidovirus helicases, an N-terminal zinc-binding domain (ZBD), stalk, and a 1B domain (Hao et al., 2017; Jia et al., 2019; Lehmann et al., 2015b). The role of these domains in nsp13 function *in vivo* is unclear, but substitutions in these domains, or the interfaces between them, have disruptive effects on *in vitro* helicase activity as well as on viral propagation (Dinten et al., 1996, 2000; Hao et al., 2017; Marle et al., 1999; Seybert et al., 2005).

Biochemical and biophysical studies suggest that nsp13 and nsp12 interact (Adedeji et al., 2012; Jia et al., 2019), but a stable complex has not been biochemically isolated or structurally characterized, and the mechanistic role of the helicase in SARS-CoV-2 replication-transcription is unknown. Here, we generate a stable nsp13:holo-RdRp:RNA complex and determine its structure by cryo-electron microscopy (cryo-EM) to 3.5 Å nominal resolution. The architecture of the complex provides constraints on models for nsp13 function and suggests a possible role for nsp13 in generating backtracked replication/transcription complexes for proofreading, template-switching during sub-genomic RNA transcription, or both. Our structure also resolves ADP-Mg^2+^ in the active site of the nidovirus-specific NiRAN domain, defining a new pocket for anti-viral therapeutic development.

## RESULTS

### The nsp13 helicase forms a stable complex with the holo-RdRp elongation complex

To investigate the potential holo-RdRp:nsp13 complex and its role in replication/transcription, we prepared active recombinant SARS-CoV-2 holo-RdRp and nsp13 (Figure S1, S2a). We used native electrophoretic mobility shift assays to test binding of holo-RdRp and nsp13 to an RNA scaffold (Figure 1a, b). Nsp12 required nsp7:8 to bind the RNA (Figure 1b, compare lanes 2 and 3), and the addition of nsp13 to the holo-RdRp:RNA complex caused a super-shift, indicating stable complex formation (Figure 1b, lane 6). Addition of ADP-AlF_3_ (Chabre, 1990), which inhibits nsp13 ATPase activity (Figure S2c), sharpened the super-shifted band (Figure 1b, lane 7), suggesting formation of a more stable, homogeneous complex. A stable complex was also detected using native mass-spectrometry (nMS), revealing a stoichiometry of nsp7:nsp8_2_:nsp12:nsp13 + RNA scaffold (Figure 1c). The ATPase activity of nsp13 was maintained within the complex (Figure S2a).

**Figure 1.**
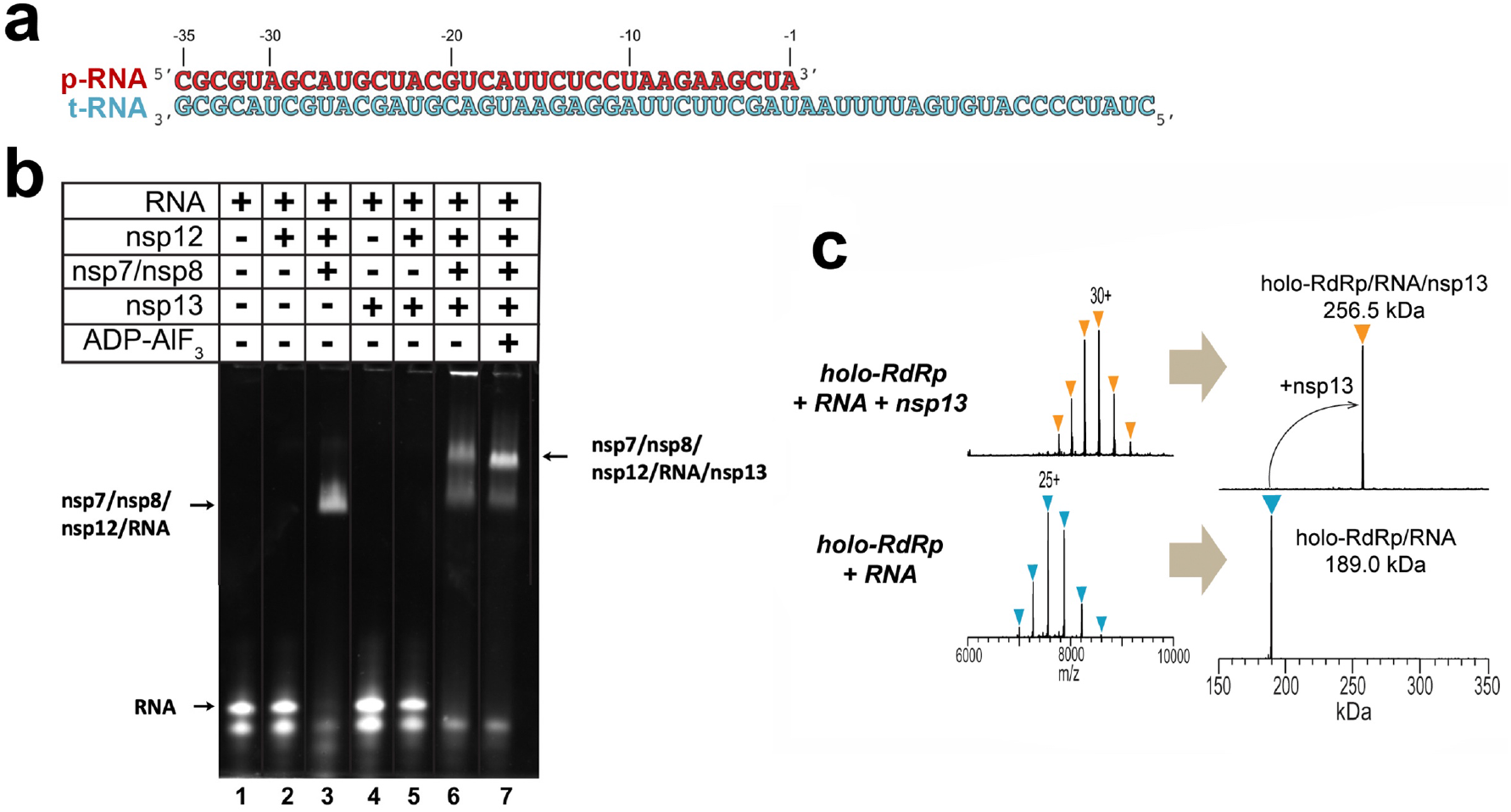
SARS-CoV-2 nsp13 helicase forms a stable complex with the holo-RdRp and an RNA scaffold. **a**. RNA scaffold used for biochemistry, native mass spectrometry (nMS), and cryo-EM. **b**. Native gel electrophoretic mobility shift assay reveals that nsp13 forms a stable complex with holo-RdRp:RNA. The 4.5% polyacrylamide gel was visualized with Gel Red to stain the RNA. **c**. nMS spectra and the corresponding deconvolved spectra for the holo-RdRp containing the RNA scaffold (a) with and without nsp13. The measured mass for the holo-RdRp:RNA complex corroborates the established stoichiometry of 1:2:1:1 for nsp7:nsp8:nsp12:RNA (Hillen et al., 2020; Kirchdoerfer and Ward, 2019; Wang et al., 2020; Yin et al., 2020), respectively (bottom). Addition of the 67.5-kDa nsp13 helicase to the RNA-bound holo-RdRp holo sample forms a transcription complex/helicase assembly with 1:1 stoichiometry (top). See also Figure S1 and S2.

### Structure of the nsp13:holo-RdRp:RNA complex

To determine the structural organization of the nsp13(ADP-AlF_3_):holo-RdRp:RNA complex (hereafter called the nsp13-replication/transcription complex, or nsp13-RTC), we analyzed the samples by single particle cryo-electron microscopy (cryo-EM). An initial dataset collected in the absence of detergent (Figure S3a) yielded poor maps due to severe particle orientation bias (Figure S3b-d). The particle orientation bias was eliminated by adding the detergent CHAPSO, previously shown to reduce preferred particle orientation by limiting particle adsorption to the air/water interface (Chen et al., 2019), yielding isotropic maps (Figure S4, S5).

The sample comprised three major classes: 67% of the particles belong to an nsp132-RTC, 20% form nsp131-RTC, and 13% form a dimer of the nsp132-RTC [(nsp132-RTC)2; Figure S4a). The major class (nsp132-RTC) yielded a 3D reconstruction and refined model at a nominal resolution of 3.5 Å (Figure 2; Figure S4b-f; Table S1). The structure resolves the complete, post-translocated holo-RdRp/RNA complex to roughly between 2.8 - 3.3 Å resolution (Figure S4b), including the nsp8 N-terminal extensions that interact with the upstream duplex RNA and the complete NiRAN domain (Hillen et al., 2020; Wang et al., 2020).

**Figure 2.**
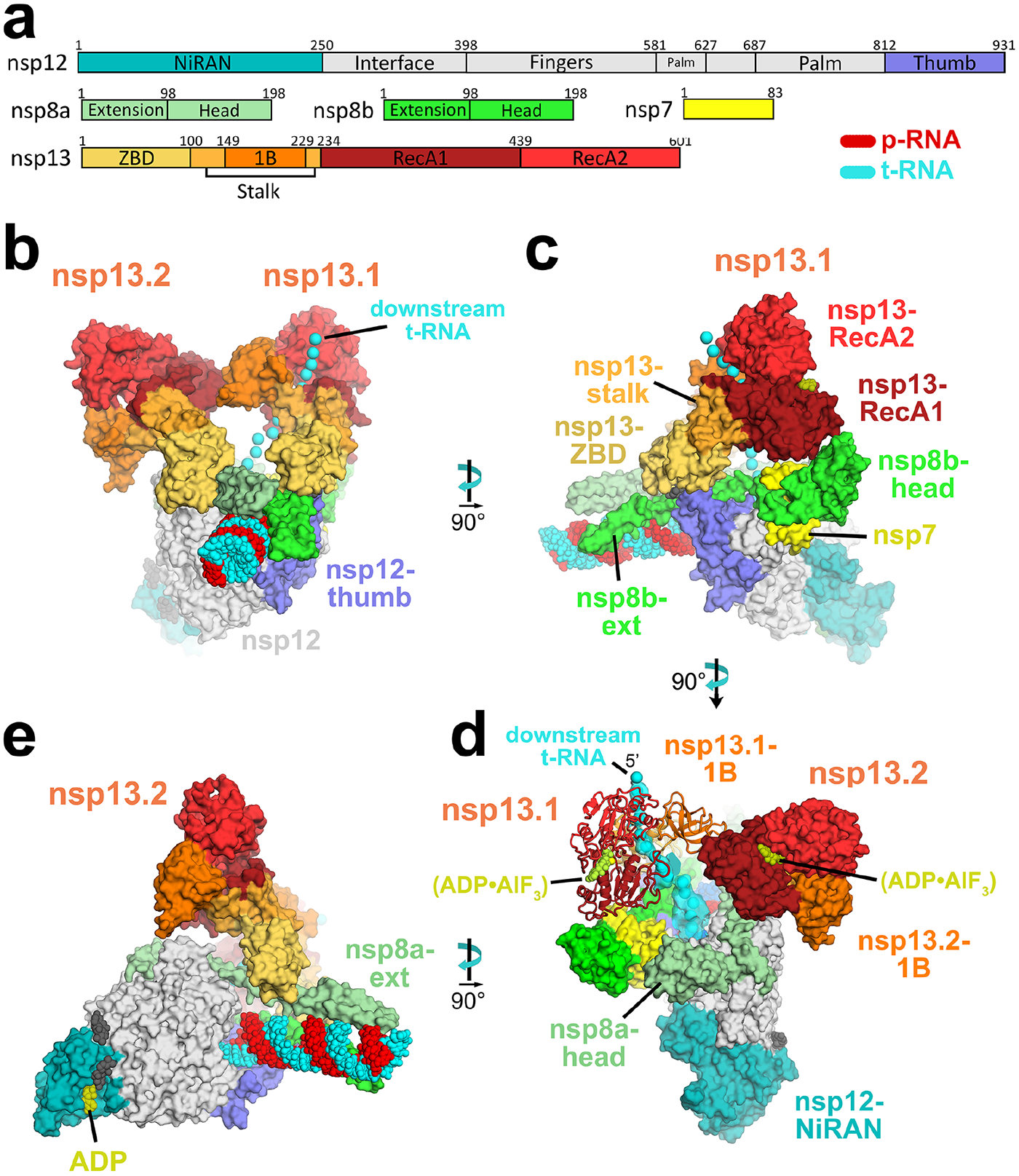
Overall structure of the SARS-CoV2 nsp13 helicase with the holo-RdRp:RNA replication/transcription complex (RTC). **a.** Schematic illustrating domain structure of SARS-CoV-2 holo-RdRp (nsp7, nsp8, nsp12) and nsp13. Structural domains discussed in the text are labeled. The color-coding corresponds to the figures throughout this manuscript unless otherwise specified. **b-e.** Orthogonal views showing the overall architecture of the nsp132-RTC. Proteins are shown as molecular surfaces (except nsp13.1 in panel **d**), RNA as atomic spheres. Adventitious CHAPSO detergent molecules are shown as atomic spheres and colored dark grey. The path of downstream t-RNA through the nsp13.1 helicase, shown as cyan spheres, is revealed with low-pass filtered (6 Å) difference density (shown in panel **d**). **b**. Two copies of the nsp13 helicase bind to the RTC. Nsp13.1 forms a tripartite interaction with the nsp8b-extension and the nsp12-thumb via the nsp13.1-ZBD. The 5’-end of the tRNA extrudes through the nucleic acid binding channel of nsp13.1. The two helicases interact via the nsp13.1-1B domain and the nsp13.2-RecA1 domain. **c.** In addition to the nsp13.1-ZBD:nsp8b-extension:nsp12-thumb tripartite interaction, nsp13.1-RecA1 interacts with nsp7 and the nsp8b-head. **d.** ADP-AlF_3_ is modeled in the NTP binding site of each helicase. the low-pass filtered (6 Å) cryo-EM difference density revealing the path of the downstream t-RNA is shown (cyan surface). **e.** The nsp13.2-ZBD interacts with the nsp8a-extension. ADP-Mg^2+^ is bound to the NiRAN domain. See also Figure S3-S5 and Table S1.

In the nsp132-RTC, the nidovirus-specific nsp13-ZBDs (Figure 2a) interact with the nsp8 N-terminal extensions; one nsp13-ZBD (nsp13.1-ZBD) interacts with nsp8b, while the other (nsp13.2-ZBD) interacts with nsp8a (Figure 2). The nsp13.1-ZBD interaction with the holo-RdRp is tripartite; besides the nsp8b the interaction incudes the tip of the nsp12-thumb domain (Figure 2c). These interactions all involve multiple residues that are universally conserved among a- and β-CoVs (Figure 3).

**Figure 3.**
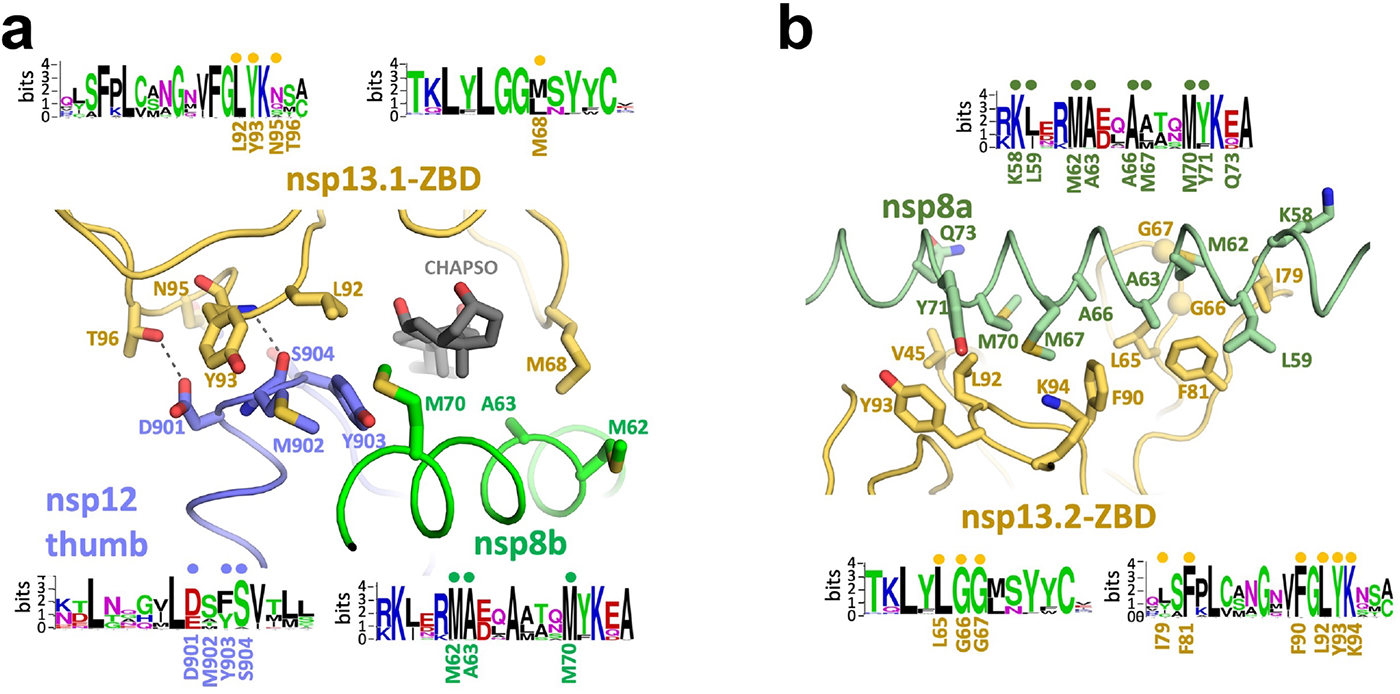
Interactions of the nsp13-ZBDs with the RTC. **a.-b.** Views of the nsp13-ABD:RTC interactions. Proteins are shown as a-carbon backbone worms. Side chains that make protein:protein interactions are shown. Polar interactions are shown as dashed grey lines. Sequence logos (Schneider and Stephens, 1990) from alignments of a- and β-CoV clades for the interacting regions are shown. The residues involved in protein:protein interactions are labeled underneath the logos. Dots above the logos denote conserved interacting residues. **a.** View of the tripartite nsp13.1-ZBD:nsp8b-extension:nsp12-thumb interaction. The adventitiously-bound CHAPSO molecule is shown in dark grey. **b.** View of the nsp13.2-ZBD:nsp8a-extension interaction. See also Figure S6.

In addition to the nsp13.1-ZBD:nsp8b:nsp12-thumb tripartite interaction, nsp13.1-RecA1 is braced against nsp7 and the nsp8b-head (Figure 2c). Other than the conserved nsp13.2-ZBD:nsp8a interaction (Figure 3b), nsp13.2 only makes additional interactions with nsp13.1 (Figure 2b). In our cryo-EM reconstructions, we note that nsp13.2 is never present without nsp13.1 (Figure S3, S4), and the disposition of nsp13.1 in nsp131-RTC is nearly identical to its disposition in nsp132-RTC (Figure S6a). Thus, nsp13.1 appears to be stably bound whereas nsp13.2 is dissociable, explaining the nMS stoichiometry (Figure 1c) and cryo-EM observations.

The holo-RdRp/RNA complex and interactions with the nsp13-ZBDs were well resolved in the cryo-EM map (Figure S4b), but structural heterogeneity in the nsp13-1B and RecA domains gave rise to poorly resolved maps that could not be improved by local refinement procedures. Maps filtered to the local resolution (Cardone et al., 2013) were used to model the overall domain orientations, and difference maps revealed that the added ADP-AlF_3_ was bound, but the low resolution of the maps prevented detailed modeling.

The nsp13.1-ZBD interface with the holo-RdRp includes an adventitious CHAPSO molecule (Figure 3a). In the absence of CHAPSO, the cryo-EM reconstructions were not suitable for detailed modeling and refinement due to severe particle orientation bias (Figure S3), but domain orientations could be modeled. Comparison with the no-detergent nsp132-RTC map revealed that the presence of CHAPSO slightly altered the disposition of the nsp13.1-ZBD with respect to nsp8b, but the overall architecture of the complex was the same (Figure S6b). The ATPase activity of nsp13 was not significantly affected by the presence of CHAPSO, whether the helicase was alone or in the RTC complex (Figure S2b). The nsp13.1-ZBD:nsp8b:nsp12-thumb, and nsp13.2-ZBD:nsp8a interfaces involve multiple residues that are universally conserved across a- and β-CoV clades (Figure 3), confirming the biological relevance of the complex.

### The NiRAN domain binds ADP-Mg^2+^

Nidovirus RdRps contain an N-terminal domain that is lacking in other viral RdRps, the NiRAN domain. The target of the NiRAN domain nucleotidylation activity is unknown, but the activity is essential for viral propagation in EAV and SARS-CoV (Lehmann et al., 2015a). In addition to ADP-AlF_3_ bound in the NTP site of each nsp13 helicase, ADP-Mg^2+^ was clearly resolved bound nsp12-NiRAN domain (Figure 4), prompting us to investigate this domain further. A search using our isolated NiRAN domain structure (nsp12 residues 1-250) on the DALI webserver (Holm and Laakso, 2016) revealed significant structural similarity with *Pseudomonas syringae* SelO [PDB 6EAC; Z-score of 10.7; (Sreelatha et al., 2018; Kirchdoerfer and Ward, 2019)}. SelO, with homologs widespread among eukaryotes and also common in prokaryotes (Dudkiewicz et al., 2012), is a pseudokinase, containing a kinase fold but lacking key catalytic residues of canonical kinases. Nevertheless, structural studies reveal that AMP-PNP binds in the SelO active site but is flipped relative to the orientation of ATP in canonical kinases. Functional studies show that SelO pseudokinases are in fact active enzymes that transfer AMP, rather than a phosphate group, to S, T, and Y residues of protein substrates [AMPylation, or adenylylation; (Sreelatha et al., 2018)]. These studies identified key SelO residues involved in catalysis, including K113 and R176 (coordinate ATP g-phosphate), E136 (stabilizes position of K113), N253 and D262 (coordinate active site Mg^2+^), and D252 (acts as catalytic base). Notably, the NiRAN domain was named after the discovery of its nucleotidylation activity (Lehmann et al., 2015a).

**Figure 4.**
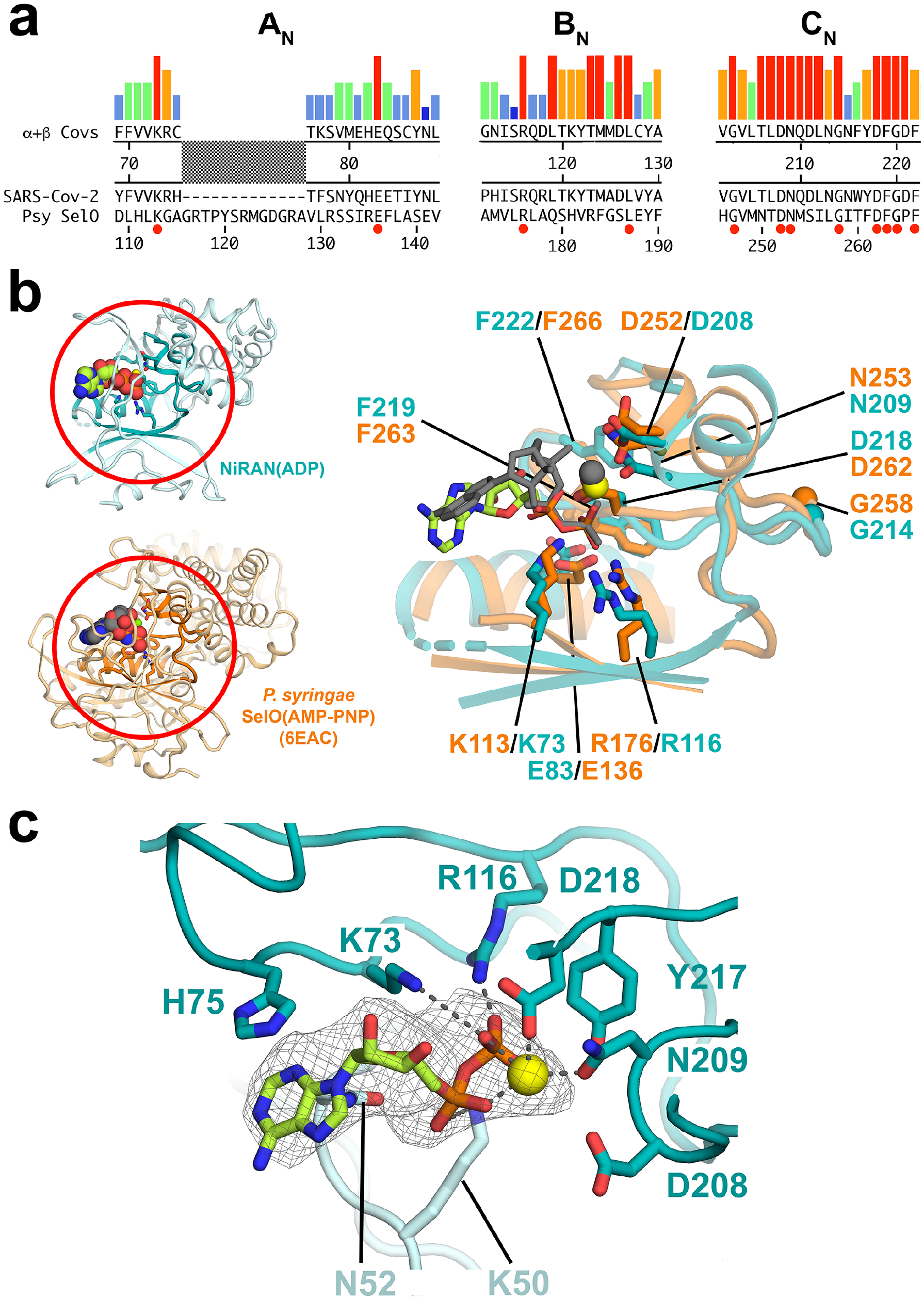
SARS-CoV-2 nsp12-NiRAN domain, pseudokinase SelO, and ADP binding. **a.** The colored histograms denote identity in a sequence alignment of 45 a- and β-CoV nsp12 sequences (red bar, 100% identity; dark blue bar, ≤ 20%) in the N-terminal signature motifs A_N_, B_N_, and C_N_ (Lehmann et al., 2015) of the NiRAN domain. The consensus sequence is shown just below. The SARS-CoV-2 nsp12 and the pseudokinase *P. syringae* (Psy) SelO (Sreelatha et al., 2018) are aligned below. Residues that are 100% identical in the nsp12 alignment and conserved in SelO are highlighted by a red dot underneath. **b.** (*left*) Structures of the SARS-CoV-2 NiRAN domain (cyan ribbon) with ADP-Mg^2+^ (spheres) and *Psy* SelO (orange) with AMP-PNP-Mg^2+^ (6EAC; Sreelatha et al., 2018). The A_N_, B_N_, and C_N_ regions are highlighted. (*right*) Structure-based alignment via a-carbons of the A_N_, B_N_, and C_N_ regions, with side chains of conserved residues shown. The a- and β-phosphates of the NiRAN-domain ADP-Mg^2+^ (limon carbon atoms, yellow sphere, respectively) superimpose almost exactly with the β- and γ-phosphates of the SelO AMP-PNP-Mg^2+^ (dark gray), whereas the nucleoside moieties diverge. **c.** The ADP-Mg^2+^-bound pocket of the SARS-CoV-2 NiRAN domain is shown. Side chains interacting with the ADP-Mg^2+^ are shown (polar interactions are denoted by gray dashed lines). D208 likely makes a water-mediated interaction with the Mg^2+^ (Sreelatha et al., 2018). The cryo-EM difference density for the ADP-Mg^2+^ is shown (light gray mesh).

Extensive sequence analysis identified three conserved sequence motifs within the nidovirus-specific N-terminal domain, A_N_, B_N_, and C_N_ [(Lehmann et al., 2015a); Figure 4a]. Overall sequence conservation between SelO and the NiRAN domain is essentially non-existent (<10%), but a structure-based alignment reveals that the key SelO catalytic residues are universally conserved within the NiRAN domain sequence motifs (Figure 4a) and are conserved in spatial organization within the aligned active-site pocket (Figure 4b).

The NiRAN domain in the nsp13-RTC structure is occupied by ADP-Mg^2+^ which is well-resolved (Figure 4c). Remarkably, the a- and β-phosphates of the NiRAN domain ADP coincide almost exactly with the β- and g-phosphates of AMP-PNP occupying the SelO active site (Figure 4b; rmsd = 0.33 Å). All of the conserved active-site residues make interactions with the ADP-Mg^2+^ as expected from the SelO structure, except NiRAN domain K73/R116 interact with the ADP β-phosphate, while the corresponding K113/R176 of SelO interact with the similarly positioned AMP-PNP g-phosphate.

### Roles of SARS-CoV-2 nsp13 in RTC processivity and backtracking

The overall architecture of the nsp13-RTC places the nucleic acid binding channel of nsp13.1 (Deng et al., 2013) directly in the path of the downstream t-RNA. Indeed, the cryo-EM difference map, low-pass filtered to 6 Å resolution, reveals the 5’-single-stranded t-RNA (Figure 1a) engaged between the RecA domains and domain 1B of nsp13.1 before entering the RdRp active site (Figure 2d). Although the interactions with nsp13.2 are not required for the formation of the nsp131-RTC (Figure S6a), nsp13.2 may stabilize the nsp13.1 complex. Biochemical analyses of nsp13 have also indicated a propensity to oligomerize, and that oligomerization can cooperatively enhance the processivity of nsp13 nucleic acid unwinding (Adedeji et al., 2012; Lee et al., 2010), suggesting a possible role for nsp13.2 in enhancing the activity and processivity of nsp13.1.

Nsp13 is an SF1B helicase, which translocate on single-stranded nucleic acid in the 5’ -> 3’ direction (Saikrishnan et al., 2009). *In vitro* studies confirm this direction of translocation for the nidovirus helicases (Adedeji et al., 2012; Bautista et al., 2002; Ivanov and Ziebuhr, 2004; Seybert et al., 2000; Seybert et al., 2000; Tanner et al., 2003). Provided the interaction of nsp13 with the holo-RdRp does not alter the unwinding polarity, which seems unlikely, the structural arrangement observed in the nsp13-RTC (Figure 2d) presents a conundrum since the RdRp translocates in the 3’ -> 5’ direction on the t-RNA strand while nsp13 would translocate on the same strand in the opposite direction. Thus, translocation of nsp13 competes with translocation by the RdRp, and if the helicase prevails it might be expected to push the RdRp backwards on the t-RNA. Reversible backwards motion of multi-subunit cellular DNA-dependent RNA polymerases (DdRps), termed backtracking, is a well-characterized feature of these enzymes (Komissarova and Kashlev, 1997b, 1997a; Nudler, 2012; Nudler et al., 1997; Wang et al., 2009).

The large active site cleft of the cellular DdRp is divided into two sections by a conserved structural element called the bridge helix (Figure 5a provides a cross-section view on the right). The main cleft on top of the bridge helix accommodates the downstream duplex DNA, while a pore or channel underneath the bridge helix forms a conduit from solution directly into the active site [Figure 5a, called the secondary channel in bacterial DdRp (Zhang et al., 1999)]. During normal transcription elongation, the NTP substrates enter the active site through the secondary channel (Westover et al., 2004; Zhang et al., 1999). In backtracking, the DdRp and associated transcription bubble move backward on the DNA, while the RNA transcript reverse threads through the complex to maintaining the register of the RNA/DNA hybrid. These movements generate a downstream segment of single-stranded RNA at the 3’-end which is extruded out the secondary channel [Figure 5a; (Abdelkareem et al., 2019; Wang et al., 2009)].

**Figure 5.**
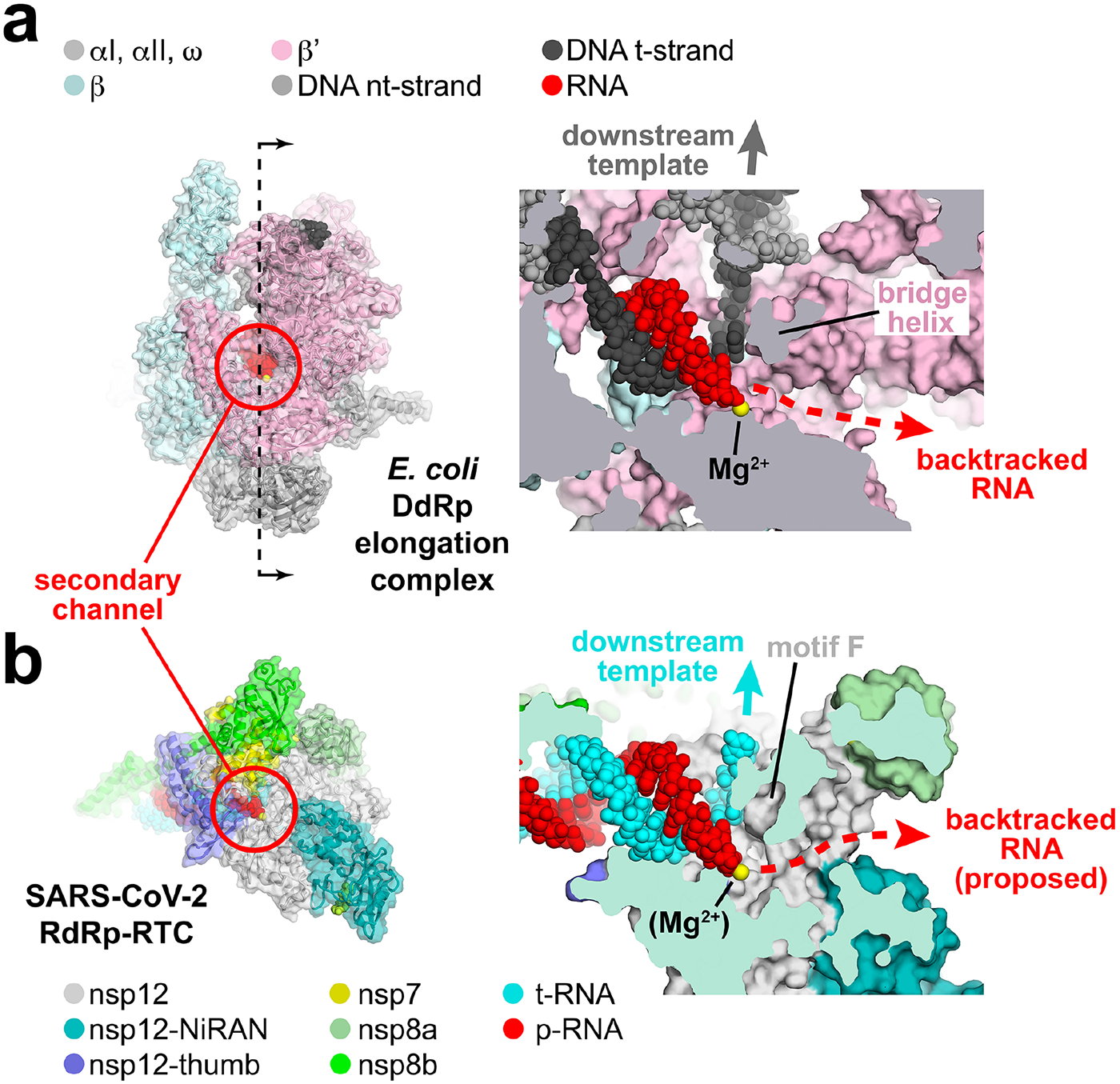
Correspondence of structural determinants for backtracking between cellular multisubunit DdRp and SARS-CoV-2 RdRp. **a.-b.** (*left*) Overall views of complexes. Proteins are shown as cartoon ribbons with transparent molecular surfaces. Nucleic acids are shown as atomic spheres. Color-coding is shown in the keys. (*right*) Cross-sectional view of the active site region (at the 3’-end of the RNA product). **a.** (*left*) Structure of the *E. coli* DdRp transcription elongation complex [EC; (Kang et al., 2017)] viewed down the secondary channel. The DdRp active site Mg^2+^-ion is shown as a yellow sphere. The secondary channel is highlighted inside the red circle. The thin dashed line illustrated the cut and viewing direction of the cross-section on the right. (*right*) Cross-sectional view showing the RNA/DNA hybrid. The bridge helix (viewed end on in cross-section) is denoted. The bridge helix directs the downstream template duplex DNA to the top (dark grey arrow). Underneath the bridge helix, the secondary channel allows NTP substrates to diffuse into the active site (Westover et al., 2004; Zhang et al., 1999) and accomodates the single-strand RNA transcript 3’-end in backtracked complexes (Abdelkareem et al., 2019; Cheung and Cramer, 2012; Wang et al., 2009). **b.** (*left*) View down the newly described secondary channel of the SARS-CoV-2 holo-RdRp. The secondary channel is highlighted inside the red circle. (*right*) Cross-sectional view showing the t-RNA/p-RNA hybrid. Motif F (viewed end on in crosssection) is denoted. Motif F directs the downstream t-RNA to the top (cyan arrow). Underneath motif F, the secondary channel could accomodate the single-strand p-RNA 3’-end in the event of backtracking.

Although evolutionarily unrelated to the DdRp, the SARS-CoV-2 RdRp active site has a similar general architecture (Figure 5b). Instead of a bridge helix, conserved motif F (SARS-CoV-2 nsp12 residues 544-555) of the viral RdRps (Bruenn, 2003), which comprises a β-hairpin loop, serves to separate the path for the t-RNA, which goes to the top, while underneath motif F is a channel that appears able to accommodate single-stranded nucleic acid. We thus propose that the SARS-CoV-2 RdRp may backtrack, generating a single-stranded RNA segment at the 3’-end that would extrude out the RdRp secondary channel (Figure 5b). We note that the active site of the RdRp is relatively open (compared to the cellular RNAPs); NTP substrates could enter directly from solution and would not need to enter through the secondary channel.

## DISCUSSION

We have established that the critical SARS-CoV-2 replication/transcription components nsp13 and holo-RdRp form a stable complex (Figure 1b, c) and determined a nominal 3.5 Å-resolution cryo-EM reconstruction (Figure 2; Figure S4). In the structure, the primary interaction determinant of the helicase with the RTC occurs between the nsp13-ZBDs and nsp8. Both of these structural elements are unique to nidoviruses, and the interaction interfaces are conserved within a- and β-CoV genera (Figure 3), indicating that this interaction represents a crucial facet of SARS-CoV-2 replication/transcription. A protein-protein interaction analysis for the SARS-CoV-1 ORFeome (which recapitulates the nsp13-RTC interactions observed in our structure) identifies nsp8 as a central hub for viral protein-protein interactions (Brunn et al., 2007). The structural architecture of nsp8a and nsp8b, with their long N-terminal helical extensions, provide a large binding surface for the association of an array of replication/transcription factors (Figure 2c, e).

Our structure reveals ADP-Mg^2+^ occupying the NiRAN domain active-site (Figure 4c), presumably because the sample was incubated with ADP-AlF_3_ prior to grid preparation. The ADP makes no base-specific interactions with the protein; NiRAN-H75 forms a cation-π interaction with the adenine base (Figure 4c), but this interaction is not expected to be strongly base-specific, and structural modeling does not suggest obvious candidates for base-specific interactions. The position corresponding to H75 in the NiRAN domain An alignment is not conserved (Figure 4a), suggesting that; i) this residue is not a determinant of base-specificity for the NiRAN domain active site, ii) that the NiRAN domain base-specificity varies among different nidoviruses, or iii) that NiRAN domains in general do not show base-specificity in their activity. The NiRAN domain of the EAV-RdRp appeared to prefer U or G for its activity (Lehmann et al., 2015a). We note that the NiRAN domain enzymatic activity is essential for viral propagation but its target is unknown (Lehmann et al., 2015a). Further experiments will be required to understand more completely the NiRAN domains activity, its preferred substrate, and its *in vivo* targets. Our results provide a structural basis for i) biochemical, biophysical, and genetic experiments to investigate these questions, and ii) provide a platform for anti-viral therapeutic development.

Our analysis comparing the viral RdRp with cellular DdRps revealed a remarkable structural similarity at the polymerase active sites - immediately downstream of each polymerase active site is a conserved structural element that divides the active site cleft into two compartments, directing the downstream nucleic acid template into one compartment and leaving a relatively open channel (the secondary channel) in the other compartment (Figure 5). In the cellular DdRps, the conserved structural element that divides the active site cleft is the bridge helix (Lane and Darst, 2010), and the secondary channel serves to allow NTP substrates to access the DdRp active site and to also accommodate the single-stranded 3’-RNA fragment generated during backtracking (Figure 5a).

In the viral RdRp, the downstream strand-separating structural element is the motif F β-hairpin loop. Unlike for multisubunit DdRps, NTP substrates could more easily access the RdRp active site by a route other than the secondary channel, but the secondary channel is perfectly positioned to accommodate backtracked RNA (Figure 5b). Based on this structural analogy, we propose that the viral RdRp can undergo backtracking and that the single-stranded 3’-RNA fragment so generated would extrude out the viral RdRp secondary channel (Figure 5b). We note that backtracking of Φ6 and poliovirus RdRps has been observed experimentally (Dulin et al., 2015, 2017).

Ignoring sequence variation, the energetics of backtracking by the cellular DdRps are close to neutral since the size of the melted transcription bubble and the length of the RNA/DNA hybrid in the active site cleft are maintained (any base pairs disrupted by backtracking are recovered somewhere else). For the SARS-CoV-2 RdRp, the arrangement of single-stranded and duplex nucleic acids during replication/transcription *in vivo* are not known, but *in vitro* the RdRp synthesizes p-RNA from a single-stranded t-RNA, resulting in a persistent upstream p-RNA/t-RNA hybrid. In this case backtracking is energetically disfavored since it only shortens the product RNA duplex without recovering duplex nucleic acids somewhere else. However, our structural analysis of the nsp13-RTC indicates that nsp13.1 can engage with the downstream single-stranded t-RNA (Figure 2d). Translocation of the helicase on this RNA strand would proceed in the 5’ -> 3’ direction, in opposition to the 3’ -> 5’ translocation of the RdRp on the same RNA strand. This aspect of helicase function would provide the NTP-dependent motor activity necessary to backtrack the RdRp. In cellular organisms, DdRp backtracking plays important roles in many processes, including the control of pausing during transcription elongation, termination, DNA repair, and fidelity (Nudler, 2012). We propose two potential roles for backtracking in SARS-CoV-2 replication/transcription: 1) fidelity, and 2) template-switching during sub-genomic transcription.

Backtracking by the cellular DdRps is favored when base pairing in the RNA/DNA hybrid is weakened by a misincorporated nucleotide in the RNA transcript (Nudler et al., 1997). Bacterial Gre-factors or eukaryotic SII interact with the backtracked transcription complex and stimulate endonucleolytic cleavage of the single-stranded 3’-RNA fragment, thus removing the misincorporated nucleotide and generating a new 3’-OH at the DdRp active site for transcription to resume (Borukhov et al., 1993; Fish and Kane, 2002; Kettenberger et al., 2003; Opalka et al., 2003). In this way, endonucleolytic transcript cleavage factors contribute to transcription fidelity *in vivo* (Erie et al., 1993; Thomas et al., 1998).

The replication and maintenance of the genomes of the CoV family of viruses, the largest known RNA genomes (Nga et al., 2011), is a hallmark of CoVs. These viruses encode a unique set of proofreading activities that are not found in other RNA viruses (Minskaia et al., 2006). This includes the N-terminal exonuclease (ExoN) domain of nsp14, which together with its co-factor nsp10 forms an important RNA proofreading machinery during viral replication (Denison et al., 2011; Smith et al., 2014). The ExoN is conserved in nidoviruses with genome sizes > 20 kb (Gorbalenya et al., 2006; Minskaia et al., 2006; Nga et al., 2011; Zirkel et al., 2011) and mutations in the ExoN lead to viruses with a mutator phenotype (Eckerle et al., 2007, 2010). The CoV murine hepatitis virus mutant lacking nsp14 ExoN activity was significantly more sensitive to remdesivir (Agostini et al., 2018). *In vitro*, nsp14 3’ -> 5’ ExoN activity can efficiently cleave a mismatched 3’-end from an RNA construct thought to mimic an RdRp misincorporation product (Bouvet et al., 2012). However, structural modeling shows that the ExoN active site would not be able to access a mismatched 3’-end in the RdRp active site, which is too narrow to accommodate the nsp10/nsp14-ExoN-domain proofreading machinery, a nearly 50 kDa protein. The nsp10/nsp14 proofreading machinery is thought to interact with the RTC complex (Subissi et al., 2014). We propose that nsp14 binds to the holo-RdRp with its ExoN active site at the mouth of the RdRp secondary channel where it would be encountered by the backtracked mismatched RNA 3’-end. Helicase-powered backtracking could thus play a role in maintaining replication/transcription fidelity by extruding the mismatched RNA 3’-end into the nsp14 ExoN active site for cleavage.

The efficiency with which the holo-RdRp can negotiate downstream obstacles to elongation is unknown. The nsp13 helicase could act in the 5’ -> 3’ direction on the t-RNA to disrupt stable RNA secondary structures of downstream RNA binding proteins (Figure 6b), both of which could be significant impediments to RNA elongation (Figure 6b). The helicase may function in this role distributively in order to avoid interfering with RdRp translocation. Alternatively, in the case of a fully duplex RNA template, the helicase could act processively to unwind the downstream duplex RNA, much like replicative helicases, such as DnaB in *Escherichia coli*, processively unwinding the DNA duplex in front of the replicative DNA polymerase (Kaplan and O’Donnell, 2002).

**Figure 6.**
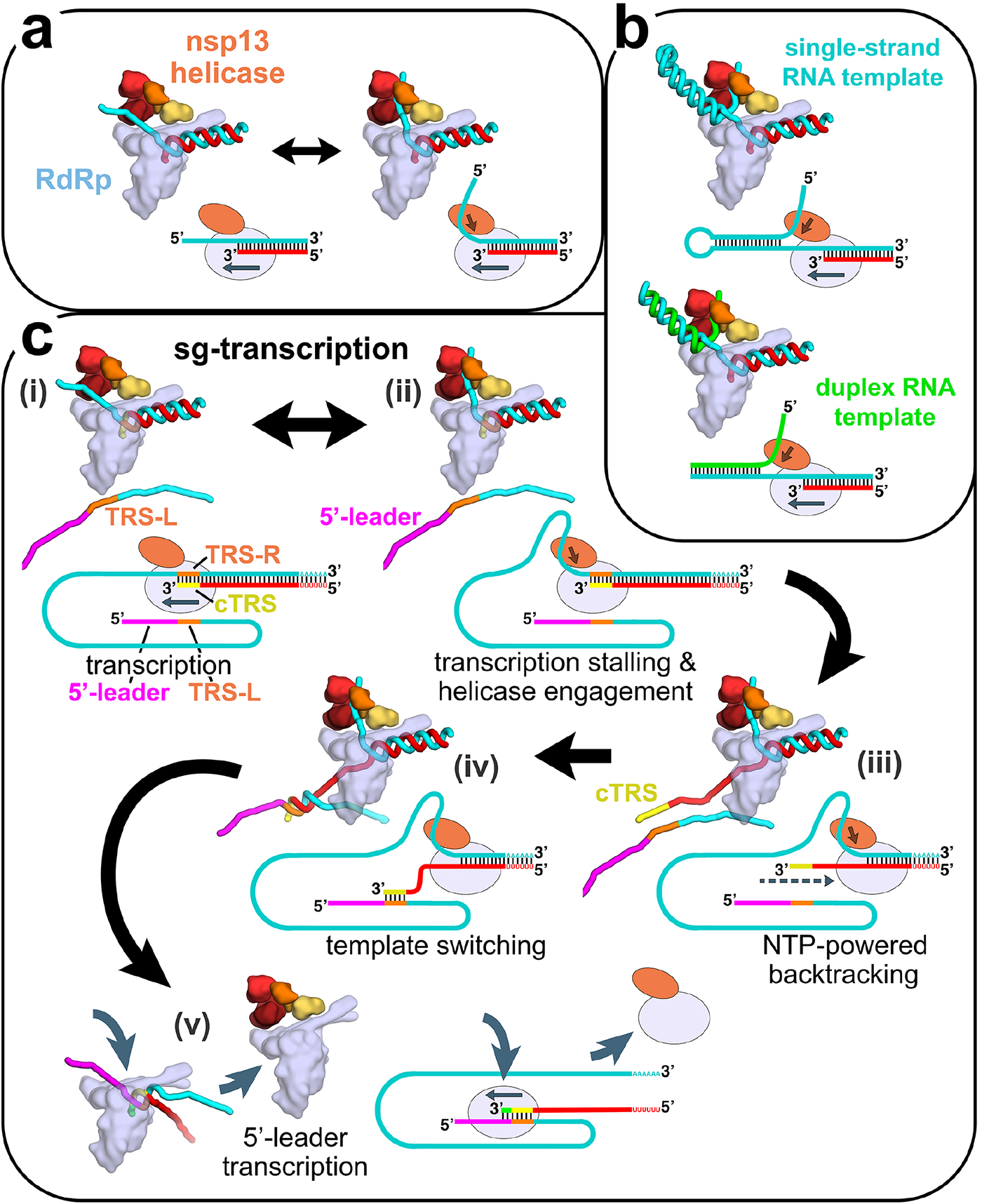
Structural basis for possible nsp13 helicase functions during viral genome replication/transcription. Structural models are shown as cartoons (holo-RdRp, light blue; nsp13.1 helicase, orange shades; RNA strands, colored tubes). The nsp13.2 helicase is not shown for clarity (all of the models are compatible with the presence of nsp13.2). With each structural diagram is a schematic cartoon illustrating the arrangement of RNA strands. Additional proteins involved in these processes are omitted. The product RNA (p-RNA) being elongated by the RdRp is shown in red. **a.** The SARS-CoV-2 nsp13-RdRp cryo-EM structure likely represents an equilibrium between two states. **b.** During RNA synthesis on a single-stranded RNA template (cyan), nsp13 could function distributively to clear downstream RNA secondary structure (or RNA binding proteins). Similarly, on a duplex RNA template (cyan and green), nsp13 could processively unwind downstream duplex RNA. **c.** Proposed helicase function during template-switching associated with sub-genomic (sg) transcription [see (Enjuanes et al., 2006; Lehmann et al., 2015b; Pasternak et al., 2001; Snijder et al., 2016; Sola et al., 2015)].

i. Negative-strand RNA synthesis proceeds from the genomic 3’-poly(A)-tail until a Transcription-Regulating Sequence [TRS-R, orange; (Alonso et al., 2002)] is transcribed (cTRS, yellow).
ii. The TRS causes transcription complex stalling.
iii. Helicase function acting on the +-strand RNA (cyan) causes backtracking of the transcription complex, freeing the pRNA 3’-end.
iv. The p-RNA 3’-end cTRS (yellow) hybridizes with the complementary TRS-L (orange) following the genomic 5’-leader sequence [magenta; (Alonso et al., 2002; Pasternak et al., 2001; Zúñiga et al., 2004)].
v. Processive helicase function backtracks the RdRp complex and unwinds the p-RNA from the genomic 3’-end. A second RdRp complex can load into the p-RNA 3’-end and continue transcription using the 5’-leader as template.

Finally, CoV transcription includes a discontinuous step during the production of sub-genomic RNAs (sg-transcription; Figure 6c) that involves a remarkable template-switching step that is a hallmark of nidoviruses (Sawicki and Sawicki, 1998). The process produces sg-RNAs that are 5’- and 3’-co-terminal with the virus genome. In this process, transcription initiates from the 3’-poly(A) tail of the +-strand RNA genome [cyan RNA in Figure 6c(i)]. Transcription proceeds until one of several transcription-regulating sequences (TRS-R, orange in Figure 6c) are encountered and transcribed (transcribed cTRS in yellow), resulting in stalling of the RTC (Alonso et al., 2002), which could allow engagement of the helicase with the t-RNA [Figure 6c(ii)]. The complement to the TRS synthesized at the 3’-end of the p-RNA (cTRS, yellow) is complementary to another TRS near the 5’-leader of the genome (TRS-L, orange; 5’-leader magenta), and the template-switching depends on complete base-pair complementarity between cTRS and TRS-L (Alonso et al., 2002; Pasternak et al., 2001; Zúñiga et al., 2004). Thus, the 6-nucleotide core-sequence of cTRS must be available to base pair with the exposed TRS-L [Figure 6d(iv)] which would be made possible by helicase-induced backtracking of the RTC to extrude the cTRL at the p-RNA 3’-end [Figure 6d(iii)]. Following hybridization of cTRS with TRS-L, a second holo-RdRp complex could load onto the ~6 base pair RNA/RNA hybrid and continue transcription of the 5’-leader [Figure 6d(v)]. This is exactly the length of the t-RNA/p-RNA duplex contacted by the nsp12 subunit of the holo-RdRp (−1 to −7, Figure 1a) and is probably what is required for a minimally stable holo-RdRp:RNA complex. Meanwhile, continued translocation of the helicase on the first RTC would processively backtrack the complex and unwind the p-RNA/t-RNA hybrid left in the wake of the RTC [Figure 6d(v)].

In addition to nidoviruses, many RNA viruses, including important pathogens such as chikungunya virus, herpesvirus, hepatitis C, Zika virus, and picornaviruses, encode their own helicases (Lehmann et al., 2015b). While these viral helicases belong to different helicase families and play different and complex roles in the viral life cycles, they all have three things in common: i) the helicases are essential for viral propagation, ii) very little is known about the precise mechanistic roles of the helicases in the viral life cycles, and iii) the helicases are thought to function as components of larger macromolecular assemblies. Our structure provides important clues to the role of the nsp13-RTC in SARS-CoV-2 replication/transcription (Figure 6) and provides guiding principles to understand the roles of the essential helicases in other RNA viruses.

## Acknowledgments

We thank A. Aher, R. Landick and C. Rice for helpful discussions, M. Ebrahim and L. Urnavicius at The Rockefeller University Evelyn Gruss Lipper Cryo-electron Microscopy Resource Center and H. Kuang at the New York Structural Biology Center (NYSBC) for help with cryo-EM data collection, T. Tuschl for support, and members of the Chait, Darst/Campbell, and Kapoor laboratories for helpful discussions. We thank B.R. tenOever for SARS-CoV-2 RNA. Some of the work reported here was conducted at the Simons Electron Microscopy Center (SEMC) and the National Resource for Automated Molecular Microscopy (NRAMM) and National Center for Cryo-EM Access and Training (NCCAT) located at the NYSBC, supported by grants from the NIH National Institute of General Medical Sciences (P41 GM103310), NYSTAR, the Simons Foundation (SF349247), the NIH Common Fund Transformative High Resolution Cryo-Electron Microscopy program (U24 GM129539) and NY State Assembly Majority. This work was supported by the Pels Family Center for Biochemistry and Structural Biology (The Rockefeller University), and NIH grants P41 GM109824 and P41 GM103314 to B.T.C, R35 GM130234 to T.K., and R35 GM118130 to S.A.D.

## Author contributions

Conceptualization; B.M., J.C., T.K, S.A.D., E.A.C. Cloning, protein purification, biochemistry; B. M., J.C., E.L., M.G., P.S., H.V. Mass spectrometry; P.D.B.O. Cryo-EM specimen preparation; B.M., E.L., J.C. Cryo-EM data collection and processing: J.C., K.M., E.E. Model building and structural analysis: S.A.D., E.A.C. Funding acquisition and supervision: B.C., T.K, S.A.D, E.A.C. Manuscript first draft: S.A.D., E.A.C. All authors contributed to finalizing the written manuscript.

## METHODS

### LEAD CONTACT AND MATERIALS AVAILABILITY

All unique/stable reagents generated in this study are available without restriction from the Lead Contact, Seth A. Darst.

### EXPERIMENTAL MODEL AND SUBJECT DETAILS

nsp7, nsp8, nsp12, and nsp13 are proteins found in the SARS-CoV-2 virus.

## METHOD DETAILS

### Protein cloning

All constructs were verified by sequencing (GeneWiz).

*Nsp7/8*. The coding sequences of the *E. coli* codon optimized SARS-CoV-2 nsp7 and nsp8 genes (gBlocks from Integrated DNA Technologies) were cloned into a pCDFDuet-1 vector (Novagen). Nsp7 bore an N-terminal His6-tag that was cleavable with preScision protease (GE Healthcare Life Sciences).

*Nsp12*. SARS-CoV-2 RNA was derived from the supernatant of propagated Vero E6 cells and provided by B.R. tenOever (Blanco-Melo et al., 2020). The sequence encoding nsp12 was reverse transcribed into cDNA using gene-specific primers and SuperScript III Reverse Transcriptase (ThermoFisher). The SARs-CoV-2 nsp12 coding sequence was subsequently cloned into a modified pRSFDuet-1 vector (Novagen) bearing an N-terminal His6-SUMO-tag which is cleavable by the ubiquitin-like protease (ULP1).

*Nsp13*. The coding sequences of the *E. coli* codon optimized SARS-CoV-2 nsp13 gene (gBlock from Integrated DNA Technologies) was cloned into a pET-28 vector (Novagen). Nsp13 bore an N-terminal His6-tag that was cleavable with preScision protease (GE Healthcare Life Sciences).

### Proetin expression and purification

#### Nsp12

The pET-SUMO plasmid expressing His6-SUMO-nsp12 was transformed into *E. coli* BL21-CodonPlus cells (Agilent) and grown overnight on LB-agar plates containing 50 μg/mL kanamycin (KAN) and 25 μg/mL chloramphenicol (CAM). Single colonies were used to inoculate LB/KAN/CAM media and grown overnight at 37°C. Two-liter flasks of LB/KAN/CAM were inoculated with 20 mL of overnight culture and ZnCl2 (10 μM final). Cells were grown at 37°C, induced at 0.6 OD600 by the addition of isopropyl β-d-1-thiogalactopyranoside (IPTG, 0.1 mM final), then incubated for 16 hr at 16°C. Cells were collected by centrifugation, resuspended in 20 mM Tris-HCl, pH 8.0, 0.1 mM ethylenediaminetetraacetic acid (EDTA)-NaOH, pH 8.0, 100 mM NaCl, 5% (v/v) glycerol, 1 mM dithiothreitol (DTT), 10 μM ZnCl2, 1x complete Protease Inhibitor Cocktail (PIC, Roche), 1 mM phenylmethylsulfonyl fluoride (PMSF), and lysed in a continuous-flow French press (Avestin). The lysate was cleared by centrifugation, then loaded onto a HiTrap Heparin HP column (GE Healthcare Life Sciences), and eluted using a gradient from 0.1 M to 1 M NaCl in TGED buffer [20 mM Tris-HCl, pH 8.0, 5% (v/v) glycerol, 0.1 mM EDTA-NaOH, pH 8.0, 1 mM DTT]. The fractions containing nsp12 were pooled and loaded onto a HisTrap HP column (GE Healthcare Life Sciences), washed, and eluted with Nickel elution buffer [20 mM Tris-HCl pH 8.0, 300 mM NaCl, 5% (v/v) glycerol, 250 mM imidazole, 1 mM 2-mercaptoethanol (BME)]. Eluted nsp12 was dialyzed overnight into dialysis buffer [20 mM Tris-HCl pH 8.0, 300 mM NaCl, 5% (v/v) glycerol, 20 mM imidazole, 1 mM BME] in the presence of His6-Ulp1 SUMO protease to cleave the His6-SUMO tag. Cleaved nsp12 was again passed through the HisTrap HP column and the flow-through was collected, concentrated by centrifugal filtration (Amicon), and loaded onto a Superdex 200 Hiload 16/600 (GE Healthcare Life Sciences) that was equilibrated with gel filtration buffer [20 mM Tris-HCl, pH 8.0, 300 mM NaCl, 5 mM MgCl2, 5% (v/v) glycerol, 2 mM DTT]. Purified nsp12 was put into storage buffer [20 mM Tris-HCl, pH 8.0, 300 mM NaCl, 5 mM MgCl2, 20% (v/v) glycerol, 2 mM DTT], aliquoted, flash frozen with liquid N2, and stored at −80°C.

#### Nsp7/8

The pCDFduet plasmid expressing His6-nsp7/8 was transformed into *E. coli* BL21 (DE3) and grown overnight on LB-agar plates containing 50 μg/mL streptomycin (Strep). Single colonies were used to inoculate LB/Strep media and grown overnight at 37°C. Two-liter flasks of LB/Strep were inoculated with 25 mL of overnight culture and ZnCl2 (10 μM final). Cells were grown at 30°C, induced at 0.6 OD600 by the addition of IPTG (0.1 mM final), then incubated for 14 hr at 16°C. Cells were collected by centrifugation, resuspended in 20 mM Tris-HCl pH 8.0, 0.1 mM EDTA-NaOH, pH 8.0, 300 mM NaCl, 5% (v/v) glycerol, 5 mM imidazole, 1 mM BME, 10 μM ZnCl2, 1x PIC (Roche), 1 mM PMSF, and lysed in a continuous-flow French press (Avestin). The lysate was cleared by centrifugation, then loaded onto a HisTrap HP column (GE Healthcare), washed, and eluted with Nickel elution buffer. Eluted nsp7/8 was dialyzed overnight into dialysis buffer in the presence of His6-Prescission Protease to cleave His6-tag. Cleaved nsp7/8 was passed through the HisTrap HP column and the flow-through was collected, concentrated by centrifugal filtration (Amicon), and loaded onto a Superdex 75 Hiload 16/600 (GE Healthcare Life Sciences) that was equilibrated with gel filtration buffer. Purified nsp7/8 was put into storage buffer, aliquoted, flash frozen with liquid N2, and stored at −80°C.

#### Nsp13

The pet28 plasmid expressing His6-nsp13 was transformed into *E. coli* Rosetta (DE3) (Novagen) and grown overnight on LB-agar plates containing 50 μg/mL KAN and 25 μg/mL CAM. Single colonies were used to inoculate LB/KAN/CAM media and grown overnight at 37°C. Two-liter flasks of LB/KAN/CAM were inoculated with 25 mL of overnight culture. Cells were grown at 37°C, induced at 0.7 OD600 by the addition of IPTG (0.2 mM final), then incubated for 17 hr at 16°C. Cells were collected by centrifugation, resuspended in 50 mM HEPES-NaOH, pH 7.5, 500 mM NaCl, 4 mM MgCl_2_, 5% (v/v) glycerol, 20 mM imidazole, 5 mM BME, 1 mM ATP, 1 mM PMSF] and lysed in a continuous-flow French press (Avestin). The lysate was cleared by centrifugation, then loaded onto a HisTrap HP column, washed, and eluted with Nickel elution buffer + 1 mM ATP. Eluted nsp13 was dialyzed overnight into 50 mM HEPES-NaOH pH 7.5, 500 mM NaCl, 4 mM MgCl_2_, 5% (v/v) glycerol, 20 mM imidazole, 5 mM BME in the presence of His6-Prescission Protease to cleave the His6-tag. Cleaved nsp13 was passed through a HisTrap HP column and the flow-through was collected, concentrated by centrifugal filtration (Amicon), and loaded onto a Superdex 200 Hiload 16/600 (GE Healthcare) that equilibrated with 25 mM HEPES-NaOH pH 7.0, 250 mM KCl, 1 mM MgCl_2_, 1 mM TCEP. Purified nsp13 was put into 25 mM HEPES-NaOH pH 7.5, 250 mM KCl, 1 mM MgCl_2_, 20% (v/v) glycerol, 1 mM TCEP, aliquoted, flash frozen with liquid N2, and stored at −80°C.

### Native electrophoretic mobility shift assays

Nsp12 (2 μM) or holo-RdRp (nsp12 incubated with three-fold molar excess nsp7/8 and additional three-fold molar excess nsp8) in transcription buffer (120 mM K-acetate, 20 mM HEPES, 10 mM MgCl_2_, 2 mM DTT) was incubated with 1 μM annealed RNA scaffold (Figure 1a) for 5 minutes at 30°C. Nsp13 (2 μM) was added where indicated. ADP and AlF_3_ (Sigma-Aldrich) was pre-mixed and added to a final concentration of 2 mM. Reaction mixtures containing nsp13 were incubated an additional 5 minutes at 30 °C. Reactions were run on a 4.5% polyacrylamide native gel (37.5:1 acrylamide:bis-acrylamide) in 1x TBE (89 mM Tris, 89 mM boric acid, 1 mM EDTA) at 4 °C. The gel was stained with Gel-Red (Biotium).

### Native mass spectrometry (nMS) analysis

The reconstituted RNA-bound holo-RdRp (RTC) and the purified nsp13 were buffer exchanged separately into nMS solution (150 mM ammonium acetate, pH 7.5, 0.01% Tween-20) using Zeba microspin desalting columns with a 40-kDa MWCO (Thermo Scientific). The buffer-exchanged samples were mixed yielding a final concentration of 4 μM RTC and 5 μM nsp13, and then incubated for 5 min at RT prior to nMS characterization.

For nMS analysis, 2–3 μL of each sample was loaded into a gold-coated quartz capillary tip that was prepared in-house and then electrosprayed into an Exactive Plus with extended mass range (EMR) instrument (Thermo Fisher Scientific) with a static direct infusion nanospray source (Olinares and Chait, 2019). The MS parameters used include: spray voltage, 1.2 kV; capillary temperature, 125 °C; in-source dissociation, 10 V; S-lens RF level, 200; resolving power, 17,500 at *m/z* of 200; AGC target, 1 x 10^6^; maximum injection time, 200 ms; number of microscans, 5; injection flatapole, 8 V; interflatapole, 4 V; bent flatapole, 4 V; high energy collision dissociation (HCD), 200 V; ultrahigh vacuum pressure, 6 – 7 × 10^-10^ mbar; total number of scans, at least 100. Mass calibration in positive EMR mode was performed using cesium iodide. For data processing, the acquired MS spectra were visualized using Thermo Xcalibur Qual Browser (v. 4.2.47). MS spectra deconvolution was performed either manually or using the software UniDec v. 4.2.0 (Marty et al., 2015; Reid et al., 2019). The deconvolved spectra were replotted using the m/z software (Proteometrics LLC). Experimental masses were reported as the average mass ± standard deviation (S.D.) across all the calculated mass values within the observed charge state series. Mass accuracies (indicated in this section in parenthesis after each measured mass) were calculated as the % difference between the measured and expected masses relative to the expected mass. For the reconstituted RTC, we obtained one predominant peak series corresponding to the fully assembled RTC with a mass of 188,984 ± 2 Da (0.008%). When nsp13 was mixed with the RTC, the resulting nMS spectrum showed a predominant peak series corresponding to the supercomplex of nsp13-RTC at 1:1 stoichiometry with a mass of 256,470 ± 10 Da (0.016%). Additional peak series were observed and were assigned to the nsp13-RTC + 490 Da (most likely from one bound Mg-ADP: 454 Da, which was also observed in the nsp13 only sample) with mass of 256,960 ± 10 Da (0.03%), and nsp12 with experimental mass of 106,787 ± 2 Da (0.0017%). The experimental masses of the individual proteins and the RNA scaffold were also determined by native MS and compared to expected masses based on their sequence and known number of zinc ions bound. The following expected masses were used for the component proteins and nucleic acid scaffold—nsp7: 9,739 Da; nsp8 (N-terminal Met lost): 21,881 Da; RNA duplex (28,681 Da); nsp13 (post-protease cleavage, has three Zn^2+^ ions coordinated with 10 deprotonated cysteine residues): 67,463 Da, and nsp12 (has two Zn^2+^ ions coordinated with 6 deprotonated cysteine residues): 106,785 Da.

### *In vitro* Transcription Assays

*In vitro* primer elongation assays were performed using a template-primer RNA scaffold of the following sequence: template (5’-CUAUCCCCAUGUGAUUUUAAUAGCUUCUUAGGAGAAUGAC-3’) and primer (5’-GUCAUUCUCCUAAGAAGCUA-3’) (Figure S1c) (Horizon Discovery Ltd./Dharmacon) annealed in 10 mM Tris-HCl, pH 8.0, 50 mM KCl, 1 mM EDTA. Primer elongation reactions (20 μL) containing 500 nM RNA scaffold, nsp12 (1μM), nsp7/8 (3μM) and NTPS (500 μM of UTP, CTP, GTP, 50 μM ATP and 1 μCi a-^32^P-ATP (Perkin-Elmer) were incubated at 30 °C for 30 min prior to quenching with an equal volume of 2x stop solution (Invitrogen-Gel Loading buffer II). Primer elongation products were separated on 15% acrylamide-6M urea denaturing gels and analyzed by phosphorimaging.

### Nsp13 helicase ATPase assays

Nsp13 ATPase activity was measured using an ATP detection method (Promega Kinase-Glo Plus^®^ kit). The assay was performed in 384-well low-volume white NBS microplates, in 20 μL volumes. Recombinant nsp13 (5 nM) and ATP (50 μM) were added to the assay buffer (20 mM HEPES, pH 7.5, 150 mM KCl, 5 mM MgCl_2_, 1 mM TCEP, 0.005% triton X100, 0.1% BSA and 2% DMSO) and incubated with gentle shaking for 10 min. Following incubation, 10 μL Kinase-Glo Plus^®^ reagent was added to the assay plate and incubated in the dark for 10 min. Relative light unit (RLU) signal was measured on a Synergy Neo2 plate reader (Biotek) in luminescent mode. For inhibition studies, nsp13 was incubated with 0-500 μM ADP:AlF_3_ (premixed in a 1:2 molar ratio) for 10 min prior to addition of ATP.

### Preparation of SARS-CoV-2 nsp13-RTC for Cryo-EM

Purified nsp12 and nsp7/8 were concentrated by centrifugal filtration (Amicon), mixed in a 1:3 molar ratio and dialyzed into 20 mM HEPES-NaOH pH 8.0, 300 mM NaCl, 10 mM MgCl_2_, 2 mM DTT, for 20 min at 22°C to remove glycerol and to allow for association. Annealed RNA scaffold (Figure 1a) was added to dialyzed nsp7/8/12 mixture and incubated for 15 min at 22°C. Sample was buffer exchanged into S6 buffer [20 mM HEPES-NaOH, pH 8.0, 150 mM K-acetate, 10 mM MgCl_2_, 2 mM DTT] using Zeba spin desalting columns. After buffer exchange, the sample was further incubated for 20 min at 30°C and then purified over a Superose 6 Increase 10/300 GL column (GE Healthcare Life Sciences) in S6 buffer. The peak corresponding to the RTC was pooled and concentrated by centrifugal filtration (Amicon). Purified nsp13 was concentrated by centrifugal filtration (Amicon) and buffer exchanged into S6 buffer using Zeba spin desalting columns, mixed with ADP and AlF_3_ (1 mM final), then added to the RTC at a molar ratio of 1:1. Complex was incubated for 5 min at 30°C.

### Cryo-EM grid preparation

Prior to grid freezing, 3-([3-cholamidopropyl]dimethylammonio)-2-hydroxy-1-propanesulfonate (CHAPSO, Anatrace) was added to the sample (8 mM final), resulting in a final complex concentration of 8 μM (Chen et al., 2019). The final buffer condition for the cryo-EM sample was 20 mM HEPES-NaOH, pH 8.0, 150 mM K-acetate, 10 mM MgCl_2_, 2 mM DTT, 1 mM ADP, 1 mM AlF_3_, 8 mM CHAPSO. C-flat holey carbon grids (CF-1.2/1.3-4Au, Protochips) were glow-discharged for 20 s prior to the application of 3.5 μL of sample. Using a Vitrobot Mark IV (Thermo Fisher Scientific), grids were blotted and plunge-froze into liquid ethane at 90% chamber humidity at 4°C.

### Cryo-EM data acquisition and processing

Structural biology software was accessed through the SBGrid consortium (Morin et al., 2013).

Grids were imaged using a 300 kV Titan Krios (Thermo Fisher Scientific) equipped with a GIF BioQuantum and K3 camera (Gatan). Images were recorded with Leginon (Suloway et al., 2005) with a pixel size of 1.1 Å/px (micrograph dimension of 5760 x 4092 px) over a defocus range of −0.8 μm to −2.5 μm and 20 eV slit. Movies were recorded in “counting mode” (native K3 camera binning 2) with ~30 e-/px/s in dose-fractionation mode with subframes of 50 ms over a 2.5 s exposure (50 frames) to give a total dose of ~66 e-/Å^2^. Dose-fractionated movies were gain-normalized, drift-corrected, summed, and dose-weighted using MotionCor2 (Zheng et al., 2017). The contrast transfer function (CTF) was estimated for each summed image using the Patch CTF module in cryoSPARC v2.15.0 (Punjani et al., 2017). Particles were picked and extracted from the dose-weighted images with box size of 256 px using cryoSPARC Blob Picker and Particle Extraction.

#### Nsp13-RTC (no detergent)

The entire dataset consisted of 10,423 motion-corrected images with 3,691,022 particles (Figure S3). Particles were sorted using three rounds of cryoSPARC 2D classification (N=100), resulting in 641,026 curated particles. An initial 3D map was generated using cryoSPARC Ab initio Reconstruction on a subset of the particles (128,437 particles from first 3,501 images). Particles were further curated using the map derived from cryoSPARC Ab initio Reconstruction as a template for cryoSPARC Heterogeneous Refinement (N=6). Class 1 consisted of 88,918 nsp8/12 particles that were further curated using cryoSPARC Ab initio Reconstruction (N=2) and refined using cryoSPARC Non-uniform Refinement. The final nsp8/12 map contained 88,918 particles with nominal resolution of 3.7 Å. Class 3 and Class 6 were each classified using Heterogeneous Refinement (N=3) revealing distinct nsp13-RTCs. Class 3.1 consisted of 29,152 (nsp132-RTC)2 particles (re-extracted with a boxsize of 320 px) that were further curated using cryoSPARC Ab initio Reconstruction (N=2) and refined using cryoSPARC Non-uniform Refinement. The final (nsp7/82/12/132/RNA)2 map consisted of 17,073 particles with nominal resolution of 4.2 Å. Class 3.3, 6.1, and 6.2 were combined and further curated with two rounds of Heterogeneous Refinement (N=3) and sorted using cryoSPARC Ab initio Reconstruction (N=3), revealing two distinct classes: nsp13-RTC and nsp132-RTC. These classes were further curated using cryoSPARC Ab initio Reconstruction (N=2) and refined using cryoSPARC Non-uniform Refinement. The final nsp131-RTC map was derived from 31,783 particles with nominal resolution of 3.8 Å. The final nsp132-RTC map was derived from 16,521 particles with nominal resolution of 4.3 Å. Local resolution calculations (Figure S3b) were generated using blocres and blocfilt from the Bsoft package (Cardone et al., 2013). The directional FSC [Figure S3c; (Tan et al., 2017)] and particle orientation distribution map from cryoSPARC (Figure S3d) for the nsp132-RTC particles indicated that the maps (and resolution esimations) were corrupted by severe particle orientation bias.

#### Nsp13-RTC (CHAPSO)

The entire dataset consisted of 4,358 motion-corrected images with 1,447,307 particles (Figure S4a). Particles were sorted using cryoSPARC 2D classification (N=100), resulting in 344,953 curated particles. Initial models (Seed 1: complex, Seed 2: decoy 1, Seed 3: decoy 2) were generated using cryoSPARC Ab initio Reconstruction on a subset of the particles (10,509 particles from first 903 images). Particles were further curated using Seeds 1-3 as 3D templates for cryoSPARC Heterogeneous Refinement (N=3), then re-extracted with a boxsize of 320 px, and followed by another round of Heterogeneous Refinement (N=3) using Seed 1 as a template. The resulting 91,058 curated particles were sorted into three classes using cryoSPARC Heterogeneous Refinement (N=3). Each class was further sorted using cryoSPARC Ab initio Reconstruction (N=3) to separate distinct 3D classes. Using these classes as references for Heterogeneous Refinement (N=6), multi-reference classification was performed on the 91,058 curated particles. Classification revealed three unique classes: (1) nsp13-RTC, (2) nsp132-RTC, (3) (nsp132-RTC)2. Particles within each class were further processed using RELION 3.1-beta Bayesian Polishing (Zivanov et al., 2018). “Polished” particles were refined using cryoSPARC Non-uniform Refinement, resulting in structures with the following particle counts and nominal resolutions: nsp13-RTC (17,345; 4.0 Å), nsp132-RTC (58.942; 3.5 Å), (nsp132-RTC)2 (11,771; 7.9 Å). Regions corresponding to nsp13.1 and nsp13.2 in the nsp132-RTC map were further refined with cryoSPARC Local Refinement. Local resolution calculations (Figure S4b) were generated using blocres and blocfilt from the Bsoft package (Cardone et al., 2013).

### Model building and refinement

For initial model of the nsp132-RTC, the initial RTC model was derived from PDB 6YYT (Hillen et al., 2020) and the initial nsp13 model from PDB 6JYT (Jia et al., 2019). The models were manually fit into the cryo-EM density maps using Chimera (Pettersen et al., 2004) and rigid-body and real-space refined using Phenix real-space-refine (Adams et al., 2010). For real-space refinement, rigid body refinement was followed by all-atom and B-factor refinement with Ramachandran and secondary structure restraints. Models were inspected and modified in Coot (Emsley and Cowtan, 2004).

### Quantification and statistical analysis

The nMS spectra were visualized using Thermo Xcalibur Qual Browser (versions 3.0.63 and 4.2.27), deconvolved using UniDec versions 3.2 and 4.1 (Marty et al., 2015; Reid et al., 2019) and plotted using the m/z software (Proteometrics LLC, New York, NY). Experimental masses (Figure 1c) were reported as the average mass ± standard deviation across all the calculated mass values obtained within the observed charge state distribution.

The local resolution of the cryo-EM maps (Figure S4b) was estimated using blocres (Cardone et al., 2013) with the following parameters: box size 15, verbose 7, sampling 1.3, and cutoff 0.5. The quantification and statistical analyses for model refinement and validation were generated using MolProbity (Chen et al., 2010) and PHENIX (Adams et al., 2010).

### Data and code availability

The cryo-EM density map for the nsp132-RTC(CHAPSO) dataset has been deposited in the EMDataBank under accession code EMD-22160. The atomic coordinates have been deposited in the Protein Data Bank under accession code 6XEZ.

## SUPPLEMENTAL INFORMATION

**Table S1.**
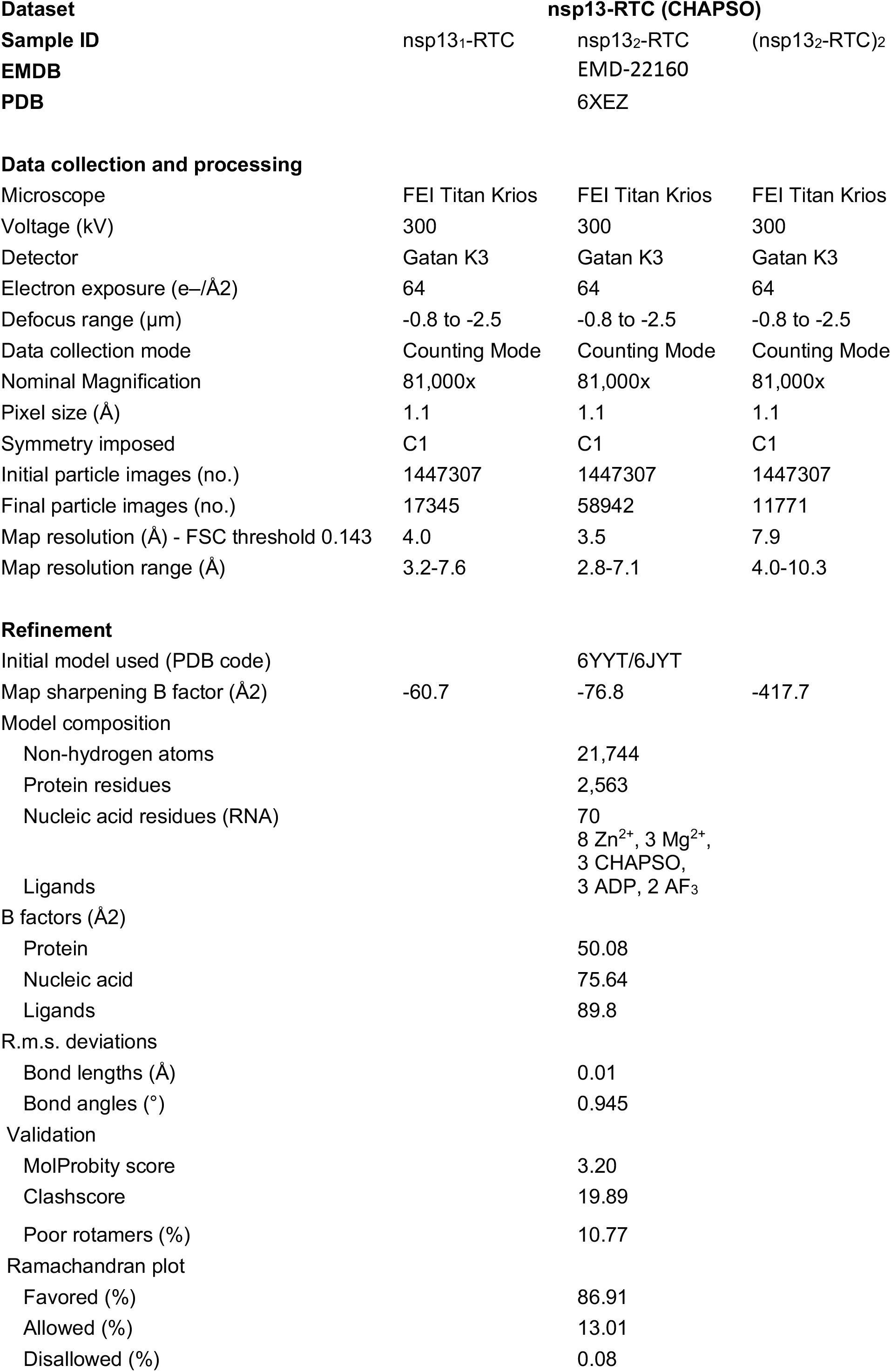
Cryo-EM data collection, refinement and validation statistics. Related to Figure 2.

**Figure S1.**
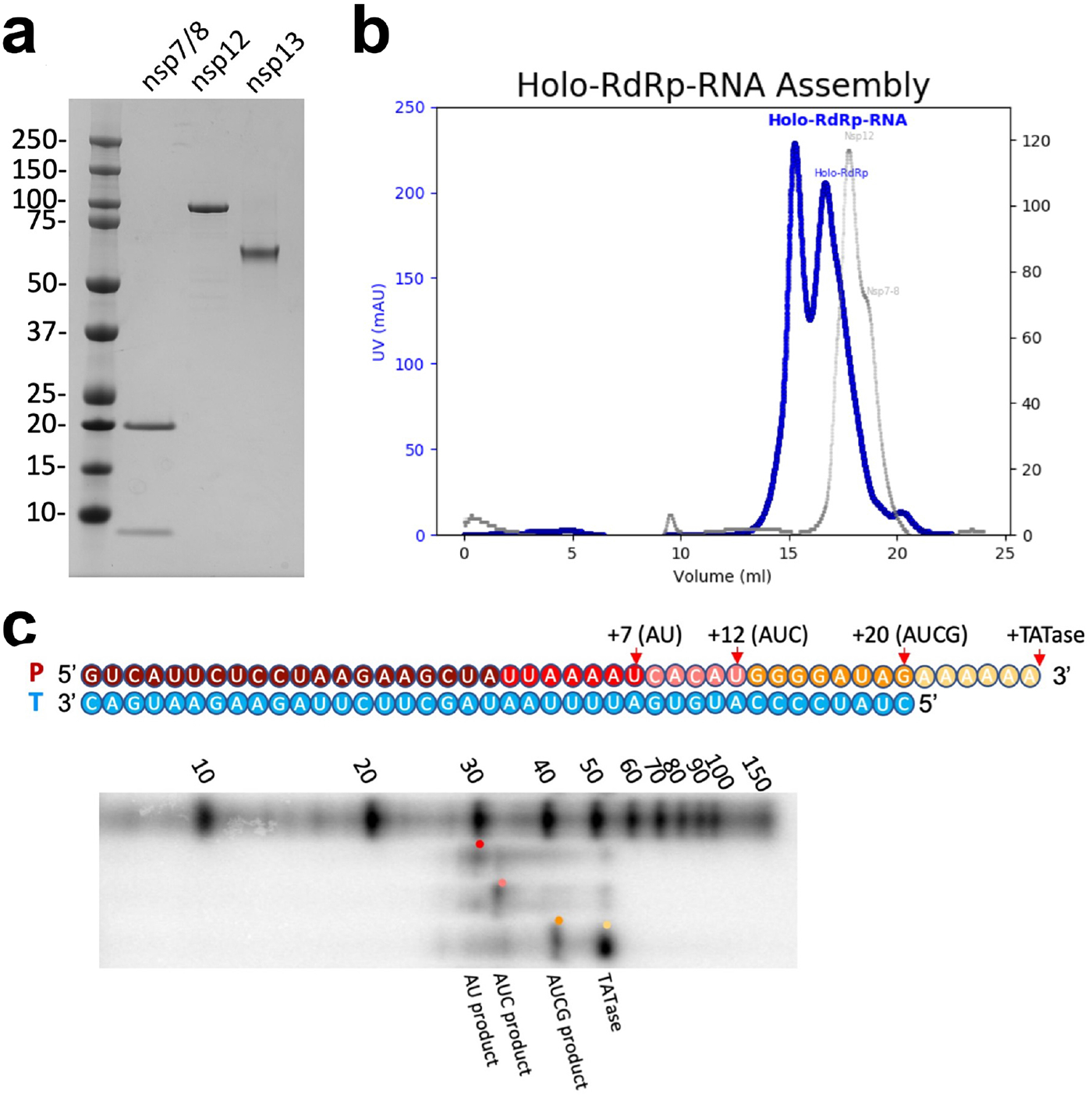
Purification and assembly of nsp13 and the RTC. Related to Figure 1. **a.** SDS-PAGE of purified SARS-CoV-2 nsp7/8, nsp12, and nsp13. **b.** Purification of RTC by size exclusion chromatography. (left) Chromatogram of RTC (blue trace) and individual components (grey trace) labeled. **c.** Holo-RdRp elongates the primer-strand of the RNA scaffold shown in the presence of NTPs. Original primer (20mer, dark red), 27mer AU elongated product (red), 32mer AUC product (pink), 40mer AUCG product (orange) and the product of nsp8-mediated TATAse activity [yellow; (Tvarogová et al., 2019). Products, taken at a 30 minute timepoint, are shown for a representative gel (n=2) and are visualized alongside Decade RNA ladder (Invitrogen).

**Figure S2.**
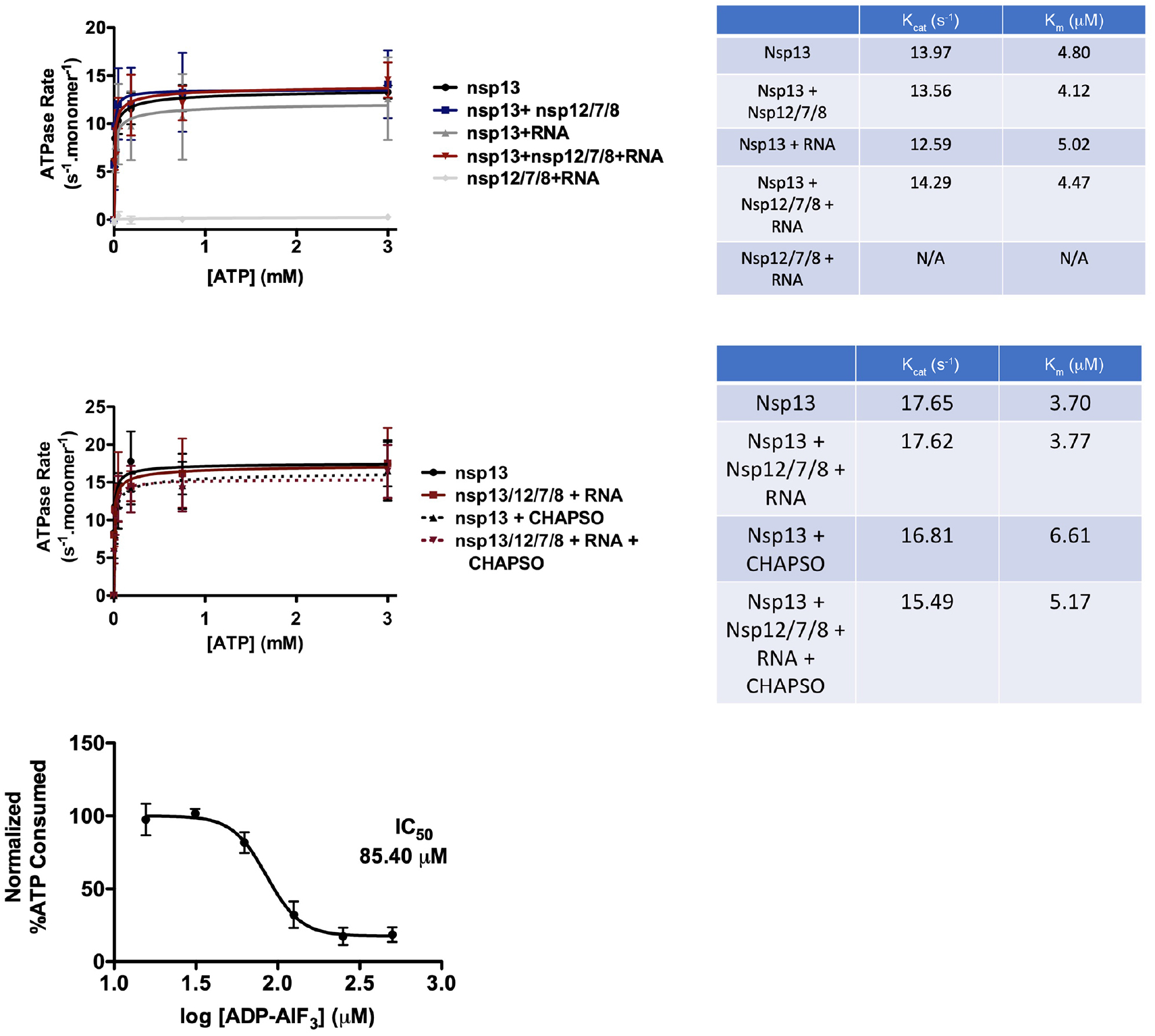
Nsp13 activities. Related to Figure 1. **a.** (*left*) ATPase assay comparing nsp13 alone, nsp13 + holo-RdRp (nsp12/7/8), nsp13 + RNA scaffold alone, nsp13-RTC (nsp13/12/7/8 + RNA), and the RTC alone (nsp12/7/8 + RNA). Error bars indicate the range for two independent measurements. (*right*) Calculated Kcat and Km values for the ATPase assay. **b.** (*left*) ATPase assay comparing nsp13 alone and nsp13-RTC (nsp13/12/7/8 + RNA) ± 8 mM CHAPSO. Error bars indicate the range for two independent measurements. (*right*) Calculated K_cat_ and K_m_ values for the ATPase assay. **c.** Inhibitory effect of ADP-AlF_3_ on nsp13 ATPase activity (N=6). Error bars denote standard deviation.

**Figure S3.**
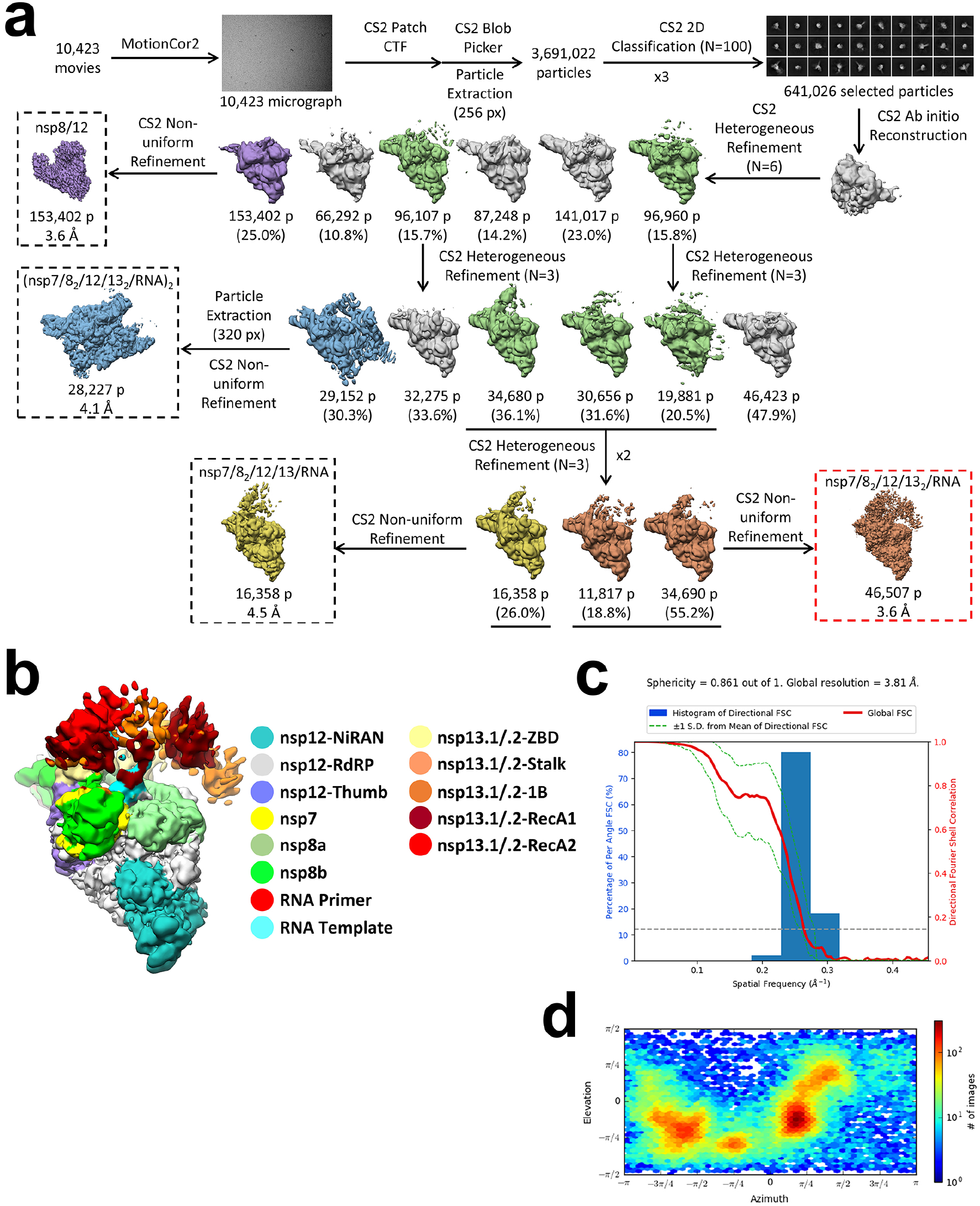
Cryo-EM processing pipeline and analysis for nsp13-RTC (no detergent) dataset. Related to Figure 2. **a.** Cryo-EM processing pipeline. **b.** Nominal 3.6 Å-resolution cryo-EM reconstruction of nsp132-RTC (no detergent) filtered by local resolution (Cardone et al., 2013) and colored by subunit according to the key on the right. **c.** Directional 3D Fourier shell correlation (FSC) for nsp132-RTC (no detergent) calculated by 3DFSC (Tan et al., 2017). **d.** Angular distribution plot for reported nsp132-RTC (no detergent) calculated in cryoSPARC. Scale shows the number of particles assigned to a particular angular bin. Blue, a low number of particles; red, a high number of particles.

**Figure S4.**
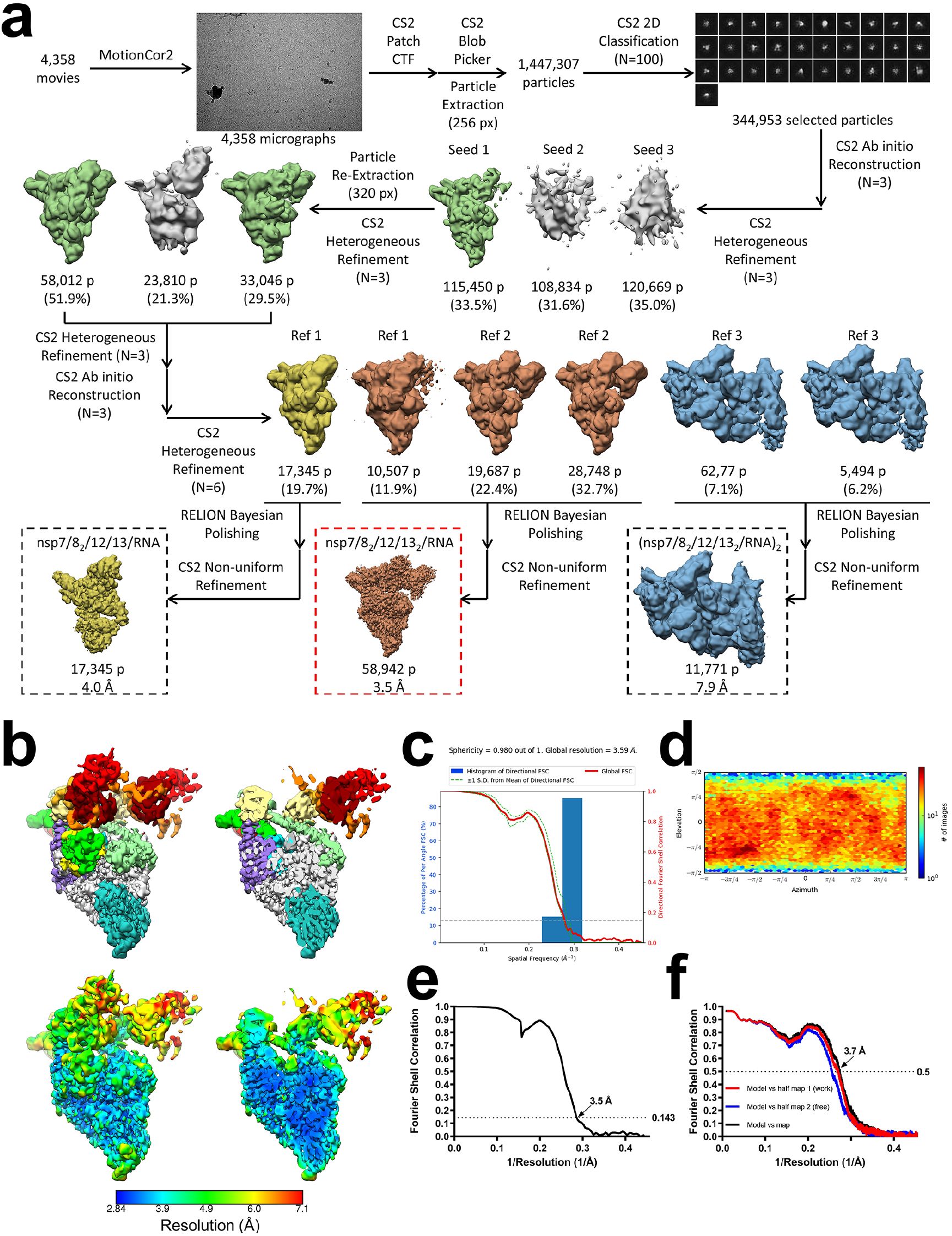
Cryo-EM processing pipeline and analysis for nsp13-RTC (CHAPSO) dataset. Related to Figure 2. **a.** Cryo-EM processing pipeline. **b.** Nominal 3.5 Å-resolution cryo-EM reconstruction of nsp132-RTC (CHAPSO) filtered by local resolution (Cardone et al., 2013). The view on the right is a cross-section. (*top*) Colored by subunit. (*bottom*) Color by local resolution (key on the bottom). **c.** Directional 3D Fourier shell correlation (FSC) for nsp132-RTC (CHAPSO) calculated by 3DFSC (Tan et al., 2017). **d.** Angular distribution plot for reported nsp132-RTC (CHAPSO) calculated in cryoSPARC. Scale shows the number of particles assigned to a particular angular bin. Blue, a low number of particles; red, a high number of particles. **e.** Gold-standard FSC plot for nsp132-RTC (CHAPSO), calculated by comparing two independently determined half-maps from cryoSPARC. The dotted line represents the 0.143 FSC cutoff which indicates a nominal resolution of 3.5 Å. **f.** FSC calculated between the refined structure and the half map used for refinement (work, red), the other half map (free, blue), and the full map (black).

**Figure S5.**
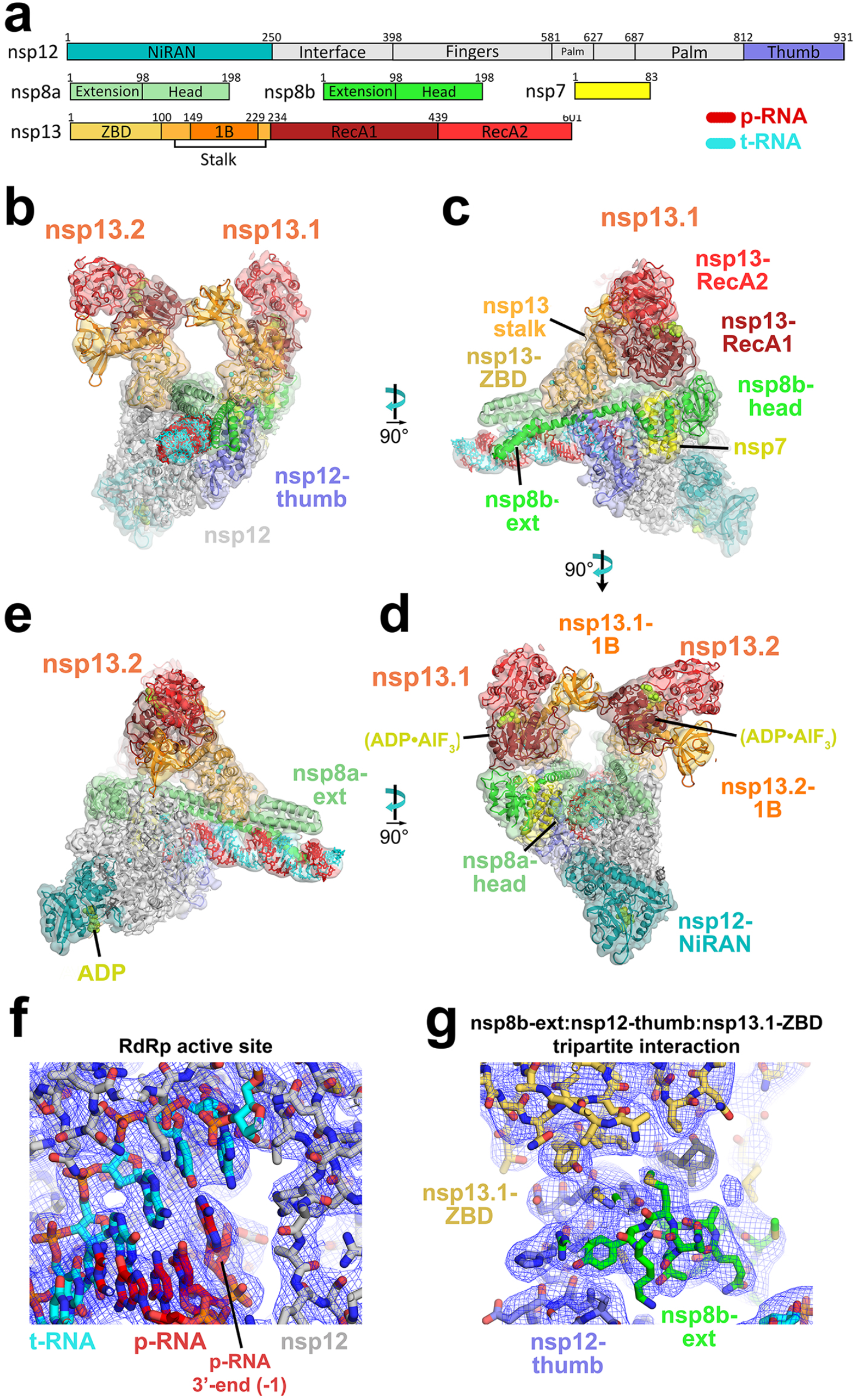
Cryo-EM density maps. Related to Figure 2. **a.** Schematic illustrating domain structure of SARS-CoV-2 holo-RdRp (nsp7, nsp8, nsp12) and nsp13. The color-coding corresponds to the figures throughout this manuscript unless otherwise specified. **b-e.** Orthogonal views showing the overall architecture of the nsp132-RTC. Shown is the transparent cryo-EM density (local-resolution filtered) with the nsp132-RTC model superimposed. Same views as Figure 2b-e. **f.** View of the nsp12 (RdRp) active site (refined model superimposed onto the cryo-EM density, shown as blue mesh), showing the post-translocated state of the RNA. **g.** View of the nsp8b-extenstion:nsp12-thumb:nsp13-ZBD tripartite interaction (refined model superimposed onto the cryo-EM density, shown as blue mesh). Similar view as Figure 3a.

**Figure S6.**
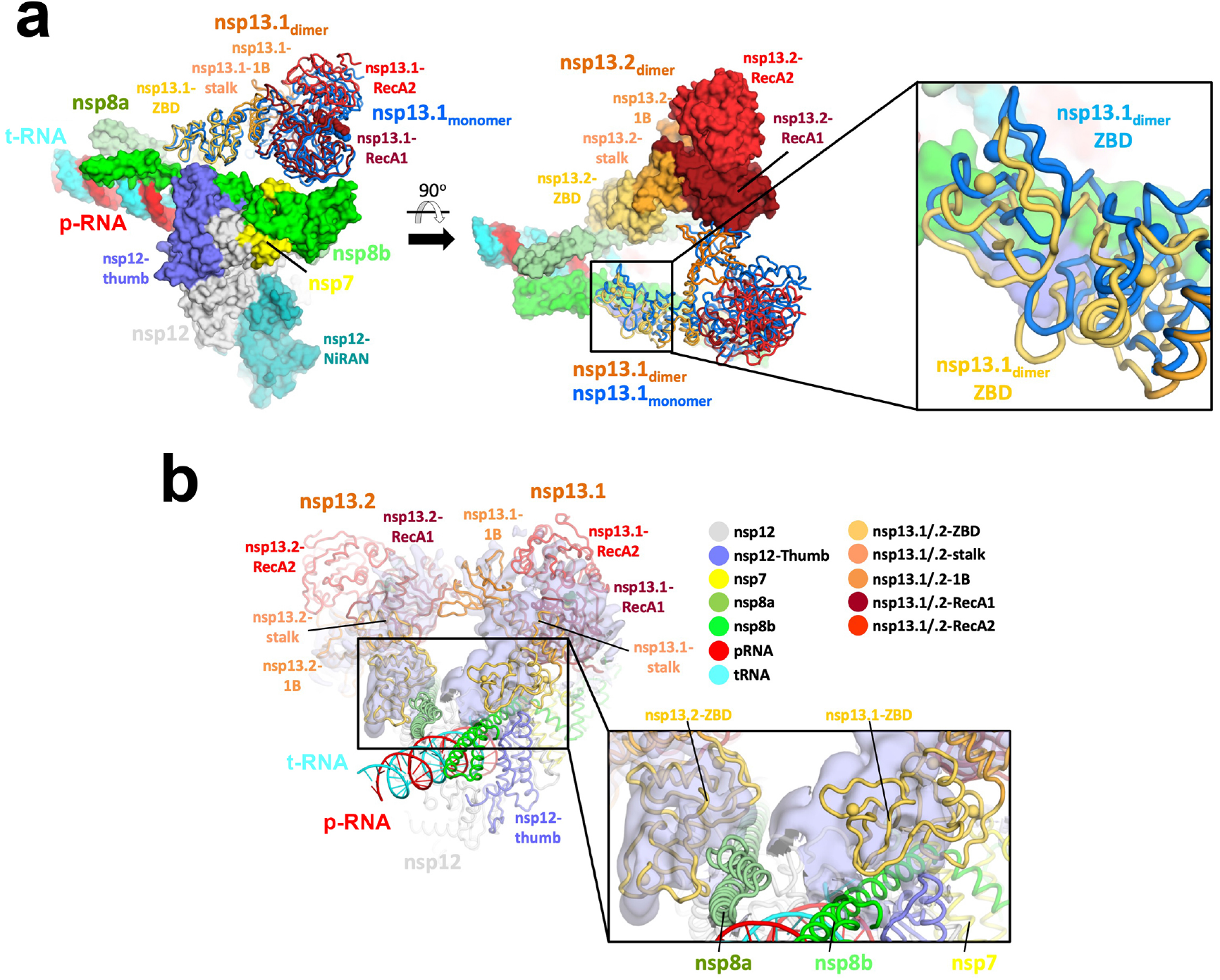
Comparison of nsp13_2_-RTC (CHAPSO) structure with nsp131-RTC (CHAPSO) and nsp132-RTC (no detergent). Related to Figure 3. **a.** Structure of nsp132-RTC (CHAPSO) colored according to key in **b** and shown as a molecular surface except nsp13.1, which is shown as cartoon tubes. Superimposed on the overall structure is nsp13 (marine) modeled from the nsp131-RTC (CHAPSO). Overall RMSD (calculated using ‘rms_cur’ in PyMOL) between the two nsp13.1 structures is 8.1 Å over 596 Cα atoms. (left) overall structure. (middle) overall structure rotated 90°. (right) zoom-in of boxed region in middle panel, showing region around nsp13.1-ZBD.s RMSD (calculated using ‘rms_cur’ in PyMOL) between the two nsp13.1 ZBDs is 3.6 Å over 100 Cα atoms. **b.** Structure of nxp132-RTC (CHAPSO) is shown in cartoon tubes, colored based on key, and superimposed onto the cryo-EM map from the nsp132-RTC (no detergent) dataset (shown as light blue transparent surface). Density map is locally filtered by resolution and difference density for nsp13 is highlighted using ‘isosurf’ command in PyMOL with 10 Å carve buffer. (left) overall structure. (right) zoom-in of boxed region in left panel, showing region around nsp13-ZBDs.

